# Enrichment of breast cancer stem cells following cytotoxic chemotherapy is mediated by hypoxia-inducible factors

**DOI:** 10.1101/2022.06.27.497729

**Authors:** Debangshu Samanta, Daniele M. Gilkes, Lisha Xiang, Pallavi Chaturvedi, Gregg L. Semenza

## Abstract

Breast cancers (BCs) that do not express the estrogen or progesterone receptor or human epidermal growth factor receptor 2 are known as triple negative breast cancers (TNBCs). Women with TNBC receive non-targeted chemotherapy with a durable response rate of less than 20%. BC stem cells (BCSCs) are a small subpopulation of BC cells that are characterized by the capacity for infinite self-renewal; are the only BC cells capable of forming a secondary (recurrent or metastatic) BC; and must be eliminated in order to eradicate BC. Hypoxia-inducible factors (HIFs) activate hundreds of genes in TNBCs and HIF-1α expression in the diagnostic tumor biopsy is associated with patient mortality. In this paper, we report that treatment of TNBC cells with cytotoxic chemotherapy increased HIF-1α and HIF-2α protein levels and HIF target gene expression. Chemotherapy also increased the percentage of BCSCs through pathways involving interleukin-6 (IL-6), IL-8, and multidrug resistance 1. HIF inhibitors blocked increased BCSC specification in response to cytotoxic chemotherapy and combination therapy led to tumor eradication. Increased HIF target gene expression in BC biopsies was correlated with increased mortality, especially in those patients treated with chemotherapy alone. Our results suggest that HIF-dependent BCSC enrichment provides a molecular and cellular basis for the high incidence of relapse in women with TNBC.

## Introduction

Over 40,000 women will die of BC in the United States this year due to the development of metastatic and chemotherapy-resistant disease. BCSCs are a subpopulation that is central to both of these processes (1, 2). Millions of BC cells enter the circulation, but only BCSCs can form a secondary (metastatic) BC (3). BCSCs are resistant to chemotherapy and the percentage of BCSCs present in the residual BC after cytotoxic chemotherapy is increased compared to the percentage of BCSCs present in the naive tumor (4–6). A reduction in BC cell burden of more than 95% may result in an apparent “complete response” to chemotherapy but may leave a population of residual BCSCs that are the source of subsequent relapse.

BC treatment is based on patient stratification into 3 groups: patients with estrogen receptor (ER) or progesterone receptor (PR)-positive BCs (60-70%), which are treated with ER antagonists or aromatase inhibitors; patients with human epidermal growth factor receptor 2 (HER2)-positive BCs (15-20%), which are treated with an anti-HER2 antibody or receptor tyrosine kinase inhibitor; and patients with TNBCs (15-20%), which do not express ER, PR, or HER2 and are treated with non-targeted chemotherapy with a durable response rate of less than 20% (7–9). Among these three groups, TNBCs are associated with the highest risk of metastatic disease and patient mortality. TNBCs are also characterized by the highest percentage of BCSCs and, at the molecular level, by a gene expression profile known as the Basal/claudin-low molecular subtype (10, 11).

Gene expression data for over 500 BCs revealed that a defining feature of the Basal/claudin-low subtype is increased expression of genes that are activated by HIFs (12). HIFs consist of an O_2_-labile HIF-1α or HIF-2α subunit and a HIF-1β subunit. HIFs function to match O_2_ delivery and O_2_ consumption, thereby preventing excessive production of reactive oxygen species (ROS) (13). ROS produced by cancer cells within tumors induce HIF activity (14). HIF-1 has been implicated in resistance to chemotherapy (15, 16) and, in hypoxic colon cancer cells, HIF-1 activated expression of multidrug resistance 1 (MDR1) (17), which mediates efflux of chemotherapy from cancer cells and is a major contributor to treatment failure (18). HIF-1α over-expression in diagnostic BC biopsies is associated with decreased patient survival in multiple studies involving thousands of BC patients (19). HIFs play multiple critical roles in the process of BC metastasis (20). Increased expression of hypoxia-induced genes in BC is also associated with poor prognosis (21). Exposure of TNBC cells to hypoxia has been shown to increase the percentage of BCSCs in a HIF-1α-dependent manner (22, 23). Treatment of cancer cells with doxorubicin was shown to induce HIF-1α expression (24). In this study, we demonstrate that when TNBCs are treated with cytotoxic chemotherapy, increased HIF activity mediates increased specification of BCSCs.

## Results

### Paclitaxel treatment increases HIF-α expression and HIF activity

MDA-MB-231 TNBC cells were exposed to 10 nM paclitaxel (Pac) for one to four days and whole cell lysates were subjected to immunoblot (IB) assays, which revealed increased HIF-1α levels on day four (Figure 1*A*). Reverse transcription-quantitative real-time PCR (RT-qPCR) assays demonstrated that that Pac increased HIF-1α mRNA expression (Figure 1*B*, left panel). Treatment of three different TNBC cell lines with Pac for four days at the drug concentration that inhibited growth by fifty percent (IC_50_; 10 nM for MDA-MB-231 and SUM-159; 5 nM for SUM-149) led to increased HIF-1α levels (Figure 1*C*). Digoxin and acriflavine are HIF inhibitors that block primary tumor growth and metastasis in mouse models of BC (25–28). Acriflavine binds to HIF-1α and HIF-2α and blocks their dimerization with HIF-1β (26), whereas digoxin blocks the accumulation of HIF-1α and HIF-2α in hypoxic cells (25). The increased HIF-1α expression induced by Pac was blocked by treatment with digoxin or acriflavine (Figure 1*D*). Pac also increased expression of HIF-2α mRNA (Figure 1*B*, right panel) and protein (Figure 1*E*), which was inhibited by co-administration of digoxin or acriflavine (Figure 1*E*). The decreased HIF-1α and HIF-2α levels in acriflavine-treated cells may result from the degradation of HIF-α subunits that have not dimerized with HIF-1β. Pac induced the expression of CA9 and ENO1 mRNA, which are products of HIF target genes, whereas expression of RPL13A mRNA, which is not HIF-regulated, was unaffected (Figure 1*F*). Pac also induced the expression of a HIF-dependent luciferase reporter gene (Figure 1*G*). The data presented in Figure 1 indicate that Pac increases HIF-1α and HIF-2α mRNA and protein expression and thereby increases HIF-dependent transcription in TNBC cells.

**Figure 1.**
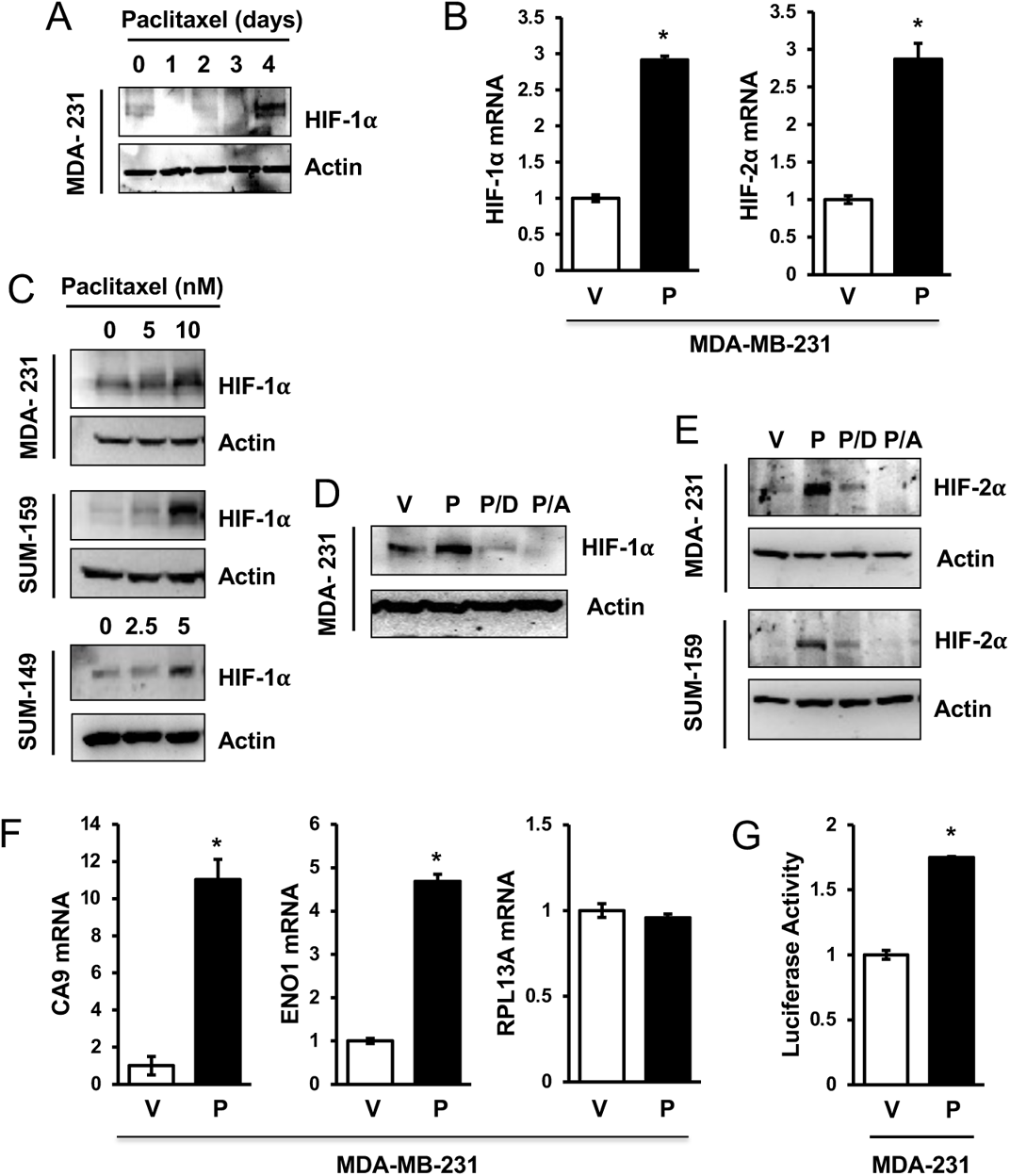
Paclitaxel induces HIF transcriptional activity. (***A***) MDA-MB-231 (MDA-231) cells were exposed to vehicle or 10 nM paclitaxel (Pac) for one to four days and cell lysates were subjected to immunoblot (IB) assays. (***B***) Cells were treated with vehicle (V) or 10 nM Pac (P) for four days and aliquots of RNA were assayed by RT-qPCR using primers specific for HIF-1α or HIF-2α mRNA relative to 18S rRNA and results were normalized to cells treated with V (mean ± SEM; *n* = 3). \**p* < 0.001 by Student’s *t* test. (***C***) Cells were treated with the indicated concentration of Pac for four days and IB assays were performed. (***D*** and ***E***) Cells were treated with vehicle (V) or 10 nM Pac, either alone (P) or in combination with 100 nM digoxin (P/D) or 1 µM acriflavine (P/A), for four days and IB assays were performed. (***F***) RT-qPCR was performed as described above (mean ± SEM; *n* = 3). \**p* < 0.001 by Student’s *t* test. (**G**) Cells were transfected with HIF-1-dependent firefly luciferase reporter p2.1 and control Renilla luciferase reporter pSV-RL, then exposed to vehicle (V) or 10 nM Pac (P) for four days, and the ratio of firefly:Renilla luciferase activity was determined. * p< 0.0001 by Student’s *t* test.

### Paclitaxel induces enrichment of cancer stem cells

To quantify BCSCs, we scored cells for aldehyde dehydrogenase 1 (ALDH) activity by flow cytometry (29, 30). BC cell lines have been shown to contain a subpopulation of ALDH^+^ cells with BCSC properties (6, 23, 30). Approximately one percent of naive TNBC cells were ALDH^+^ (Figure 2A-B). Pac treatment for four days increased ALDH^+^ cells by more than 10-fold among the surviving cells. Treatment with digoxin decreased the enrichment of ALDH^+^ cells that was induced by Pac (Figure 2A-B). BCSCs generate multicellular spheroids in suspension culture known as mammospheres (31, 32). Treatment of TNBC cells with Pac for four days enriched for mammosphere forming-cells and this induction was blocked by digoxin (Figure 2C-D). When the primary mammospheres are dissociated and re-plated, they give rise to secondary mammospheres, which reflect the self-renewal capacity of BCSCs. Treatment with digoxin alone decreased primary and secondary mammosphere formation and blocked the increased mammosphere formation that was induced by Pac (Figure 2*E*). The data presented in Figure 2 indicate that exposure of TNBCs to Pac induces BCSC enrichment that can be blocked by a HIF inhibitor.

**Figure 2.**
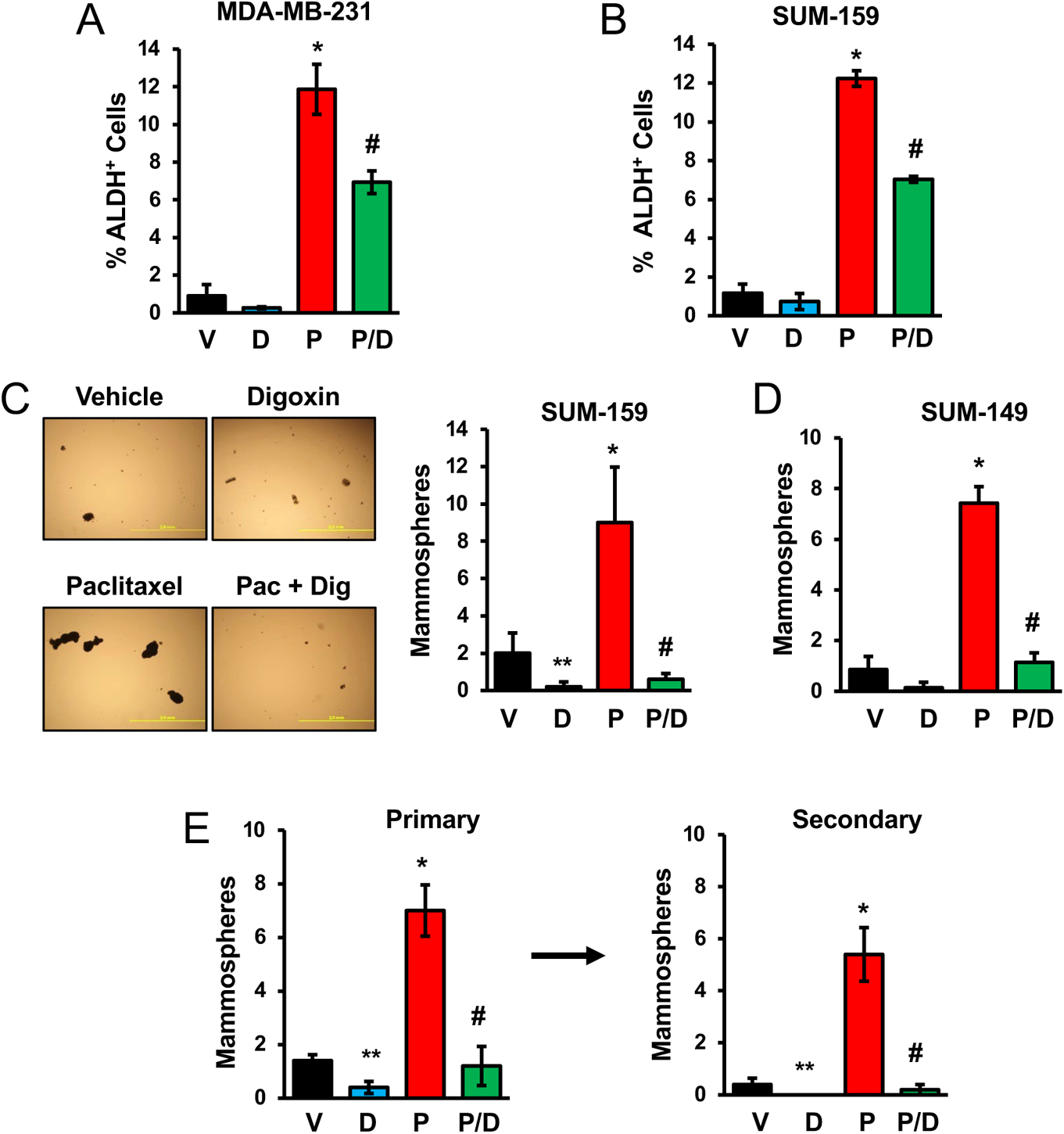
HIF inhibitors block enrichment of BCSCs by Pac. (***A*** and ***B***) Cells were treated with vehicle (V), 100 nM digoxin (D), 10 nM Pac (P), or digoxin and Pac (P/D) for four days, ALDH^+^ cells were determined by flow cytometry (mean ± SEM; n=3). \**p* < 0.001 compared to V, and ^#^*p* < 0.01 compared to P. (***C*** and ***D***) Cells were treated as described above. After four days, the cells were transferred to ultra-low attachment plates, and seven days later the number of mammospheres per field was counted (mean ± SEM; *n* = 3). \**p* < 0.001, \*\**p* < 0.01 compared to V, and ^#^*p* < 0.001 compared to P. Representative photomicrographs of mammospheres are shown; scale bar: 2 mm. (***E***) SUM-159 cells were treated as described above. Primary mammospheres were counted, collected, dissociated, transferred to ultra-low attachment plates, and after seven days secondary mammospheres were counted (mean ± SEM; n = 3). * *p* < 0.001, \*\**p* < 0.01 compared to V, and ^#^*p* < 0.001 compared to P.

### Paclitaxel increases interleukin expression

IL-6 and IL-8 promote the survival and self-renewal of BCSCs (6, 30, 33–37). Exposure of TNBC cells to Pac for four days increased IL6 and IL-8 mRNA levels, and this effect was inhibited by treatment with the HIF inhibitor digoxin or acriflavine (Figure 3A-B). Expression of RPL13A mRNA was not affected by exposure of TNBC cells to Pac, either alone or in combination with a HIF inhibitor (Figure S1). IB assays revealed that treatment of TNBCs with Pac induced increased IL-6 protein levels (Figure 3*C*), which was similar to the induction of HIF-1α in the same cell lysates (Figure 1*C*). The increased IL-6 protein expression that was induced by Pac was inhibited by concomitant treatment with digoxin or acriflavine (Figure 3*D*). Treatment with neutralizing antibodies against IL-6 or IL-8 during the exposure of TNBC cells to Pac blocked the enrichment of mammosphere-forming (Figure 3*E*) and ALDH^+^ (Figure 3*F*) BCSCs, indicating a requirement for interleukin signaling.

**Figure 3.**
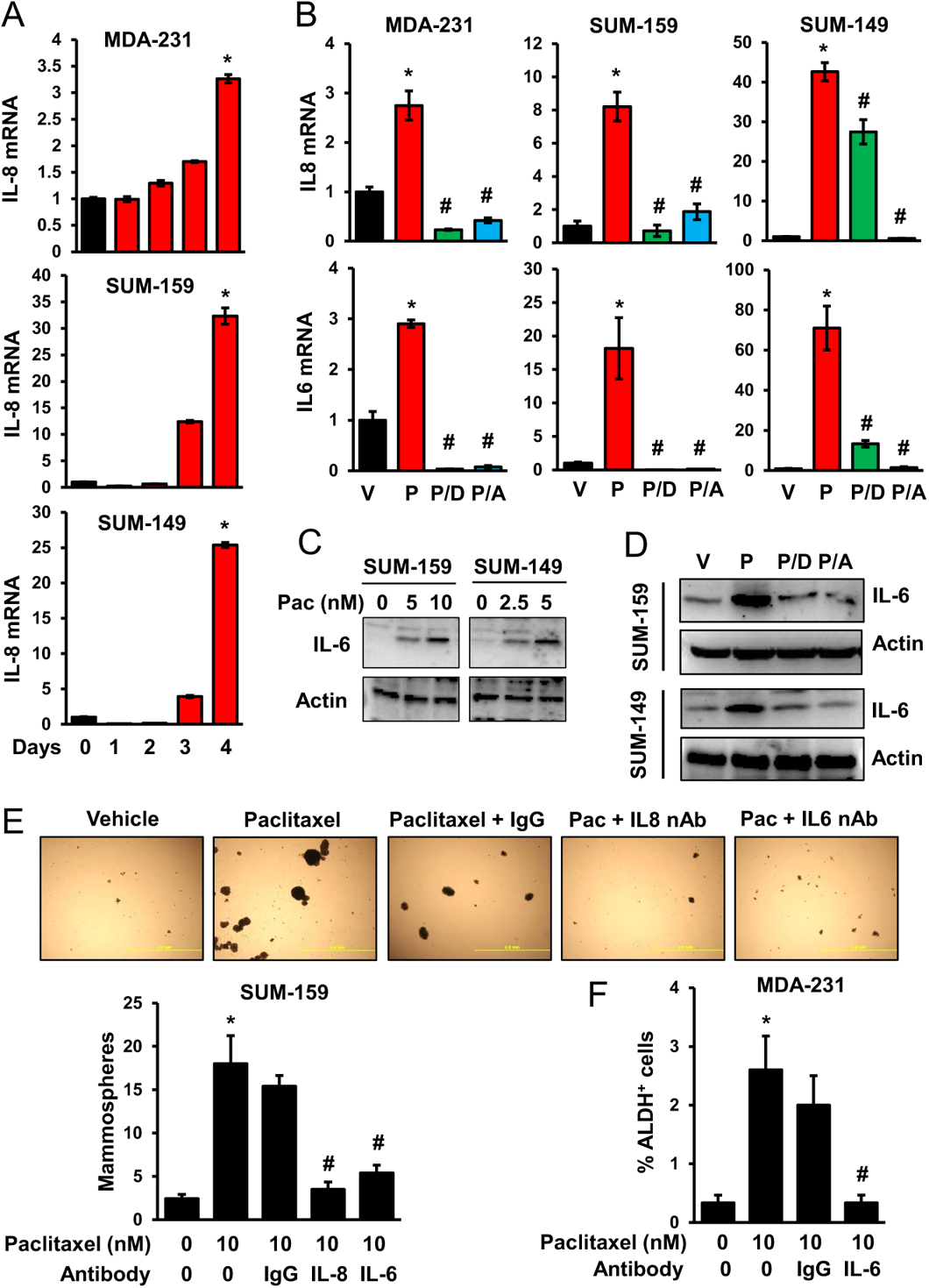
Pac increases IL-6 and IL-8 expression, leading to BCSC enrichment. (***A***) Cells were treated with 10 nM (MDA-MB-231 and SUM-159) or 5 nM (SUM-149) Pac for up to four days and RT-qPCR was performed (mean ± SEM; *n* = 3). \**p* < 0.001 compared to vehicle. (***B***) Cells were treated for four days with vehicle (V) or Pac as described above, either alone (P) or in combination with 100 nM digoxin (P/D) or 1 µM Acriflavine (P/A), and RT-qPCR was performed (mean ± SEM; *n* = 3). \**p* < 0.001 compared to V, and ^#^*p* < 0.001 compared to P. (***C***) Cells were treated with Pac for four days and IB assays were performed. (***D***) Cells were exposed to vehicle (V) or Pac, either alone (P) or in combination with digoxin (P/D) or acriflavine (P/A) for four days and IB assays were performed. (***E*** and ***F***) Cells were treated with either vehicle (V) or Pac, either alone or in the presence of IgG or neutralizing antibody (500 ng/ml) against IL-6 or IL-8 for four days, followed by mammosphere (***E***) or ALDH (***F***) assays (mean ± SEM; n = 3). \**p* < 0.001 compared to 0 nM Pac, and ^#^*p* < 0.001 compared to 10 nM Pac.

### Effect of HIF knockdown on BCSC specification induced by paclitaxel

To complement the studies described above using HIF inhibitors, we analyzed subclones of MDA-MB-231, which were stably transfected with a lentiviral vector encoding a non-targeting control (NTC) short hairpin RNA (shRNA) or shRNAs targeting HIF-1α and HIF-2α, such that there was a double knockdown [DKD] of HIF activity (28). Treatment of NTC cells with Pac increased HIF-1α, HIF-2α, and IL-6 expression and increased the percentage of ALDH^+^ cells, whereas these responses were all eliminated in DKD cells (Figure 4A-C). Analysis of subclones with knockdown of either HIF-1α or HIF-2α demonstrated that both proteins were required for maximal Pac-induced BCSC enrichment (Figure 4*D*). Pac also increased mammosphere formation in NTC cells and this response was significantly decreased in the single and double knockdown subclones (Figure 4*E*). Knockdown of either HIF-1α or HIF-2α inhibited Pac-induced IL-8 and IL-6 expression, and knockdown of both HIF-1α or HIF-2α completely blocked the response to Pac (Figure 4F-G). Thus, inhibition of HIF activity using either a pharmacologic (Figures 2 and 3) or genetic (Figure 4) approach blocked interleukin expression and BCSC enrichment that was induced by Pac.

**Figure 4.**
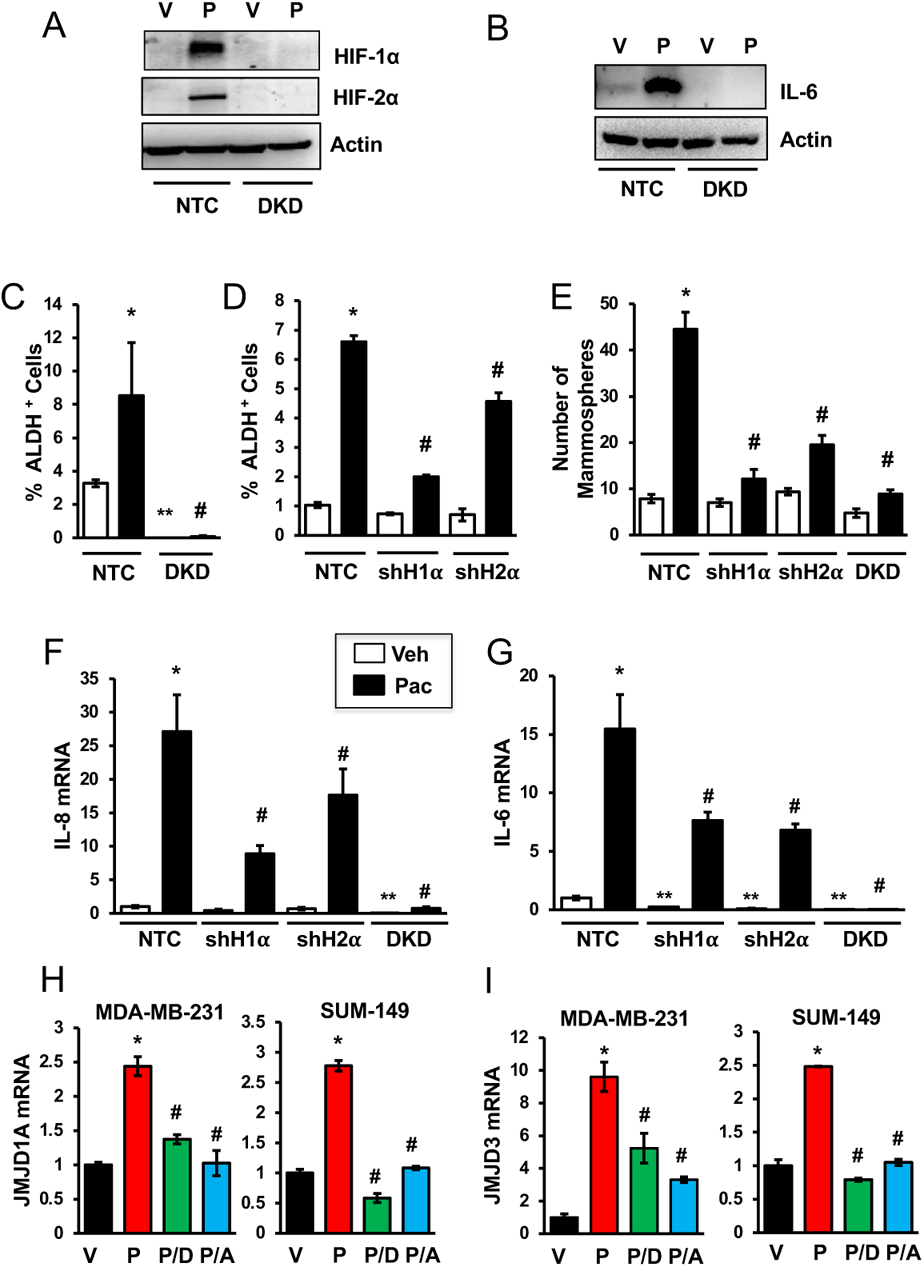
HIFs mediate increased levels of interleukins and BCSCs in response to Pac. (***A-C***) MDA-MB-231 subclones transfected with a vector encoding a non-targeting control shRNA (NTC) or vectors encoding shRNAs against HIF-1α and HIF-2α (DKD), were treated with either vehicle (V) or 10 nM Pac (P) for four days and IB (***A*** and ***B***) or ALDH (***C***) assays were performed (mean ± SEM; n=3). \**p* < 0.001 compared to vehicle-treated NTC and ^#^*p* < 0.001 compared to Pac-treated NTC. (***D***) NTC and subclones stably transfected with vector encoding an shRNA against HIF-1α (shH1α) or HIF-2α (shH2α) were treated with Pac and ALDH assays were performed (mean ± SEM; *n* = 3). \**p* < 0.001 compared to vehicle-treated NTC, and ^#^*p* < 0.001 compared to Pac-treated NTC. (**E**) Subclones were treated with Pac and mammosphere assays were performed (mean ± SEM; *n* = 3). \**p* < 0.001 compared to vehicle-treated NTC, and ^#^*p* < 0.001 compared to Pac-treated NTC. (***F*** and ***G***) Subclones were treated with V or P and analyzed by RT-qPCR (mean ± SEM; *n* = 3). \**p* < 0.001, ** *p* < 0.01 compared to vehicle-treated NTC, and ^#^*p* < 0.001 compared to Pac-treated NTC. (***H*** and ***I***) Cells were treated with vehicle (V) or Pac, either alone (P) or in combination with digoxin (P/D) or acriflavine (P/A), and RT-qPCR was performed (mean ± SEM; *n* = 3). \**p* < 0.001 compared to V, and ^#^*p* < 0.001 compared to P.

The histone demethylase JMJD1A is encoded by a HIF-regulated gene, binds to promoter sequences of the *IL8* gene and increases *IL-8* transcription (38). Treatment with Pac induced JMJD1A expression, whereas digoxin or acriflavine abrogated this effect (Figure 4H). Similarly, the histone demethylase JMJD3 is a HIF target gene product that activates *IL6* gene transcription (39). Pac treatment increased JMJD3 expression, whereas digoxin or acriflavine blocked this effect (Figure 4I). These data suggest that HIFs contribute indirectly to increased *IL8* and *IL6* expression by activating *JMJD1A* and *JMJD3* gene expression, respectively.

### Paclitaxel-induced SMAD2 and STAT3 do not increase BCSCs in the absence of HIF activity

A recent publication reported that in response to paclitaxel, TGF-β-induced SMAD2/4 expression led to increased IL-8 signaling, which was required for BCSC enrichment (6). We found that treatment with Pac increased SMAD2 phosphorylation in both NTC and DKD cells (Figure S2*A*), despite the loss of ALDH^+^ cells in the DKD subclone (Figure 4*C*). JAK2-dependent phosphorylation of STAT3 has also been implicated in BCSC specification in TNBCs (36, 40). Treatment of MDA-MB-231 subclones with Pac induced STAT3 phosphorylation in both NTC and DKD subclones (Figure S2*B*). Thus, the data presented in Figure S2 indicate that activation of SMAD2 or STAT3 in response to Pac is HIF-independent but does not lead to increased BCSCs in HIF-deficient cells.

### Enrichment of BCSCs by paclitaxel is not associated with increased TLR-4 or NF-κB

Previous reports indicated that Pac induces TLR-4 expression, which leads to increased NF-κB activity (41). However, treatment of TNBC cells with Pac at concentrations that lead to BCSC enrichment failed to increase TLR-4 mRNA levels (Figure S3A-B). Phosphorylation of IκBα was not affected by Pac treatment in MDA-MB-231, SUM-159 or SUM-149 cells (Figure S3*C*). Our results are consistent with published data indicating TNBC cells are characterized by constitutive activation of NF-κB (42). Thus, induction of TLR-4-dependent NF-κB signaling does not appear to play a major role in Pac-induced BCSC enrichment, although constitutive NF-κB activity may contribute to BCSC specification in TNBC.

### Increased ROS levels are required for HIF-1α induction and BCSC enrichment

ROS levels were increased when cancer cells were exposed to high concentrations of Pac for short periods of time (100 nM for 8 hours) (43, 44). We treated SUM-149 or SUM-159 cells with Pac at IC_50_ for four days and stained the cells with MitoSOX Red, which is targeted to mitochondria and produces fluorescence when oxidized by superoxide radicals. Pac treatment induced a significant increase in MitoSOX^+^ TNBC cells as determined by flow cytometry (Figure 5A-B). However, manganese (III) tetrakis (1-methyl-4-pyridyl) porphyrin (MnTMPyP), a cell-permeable superoxide radical scavenger, inhibited the increase in ROS by Pac (Figure 5A-B). MnTMPyP also inhibited HIF-1α expression that was induced by Pac (Figure 5C-D) and blocked the enrichment of mammosphere-forming cells (Figure 5E-F). The results presented in Figure 5 indicate that Pac increases ROS levels, which trigger increased HIF-1α expression, leading to BCSC enrichment.

**Figure 5.**
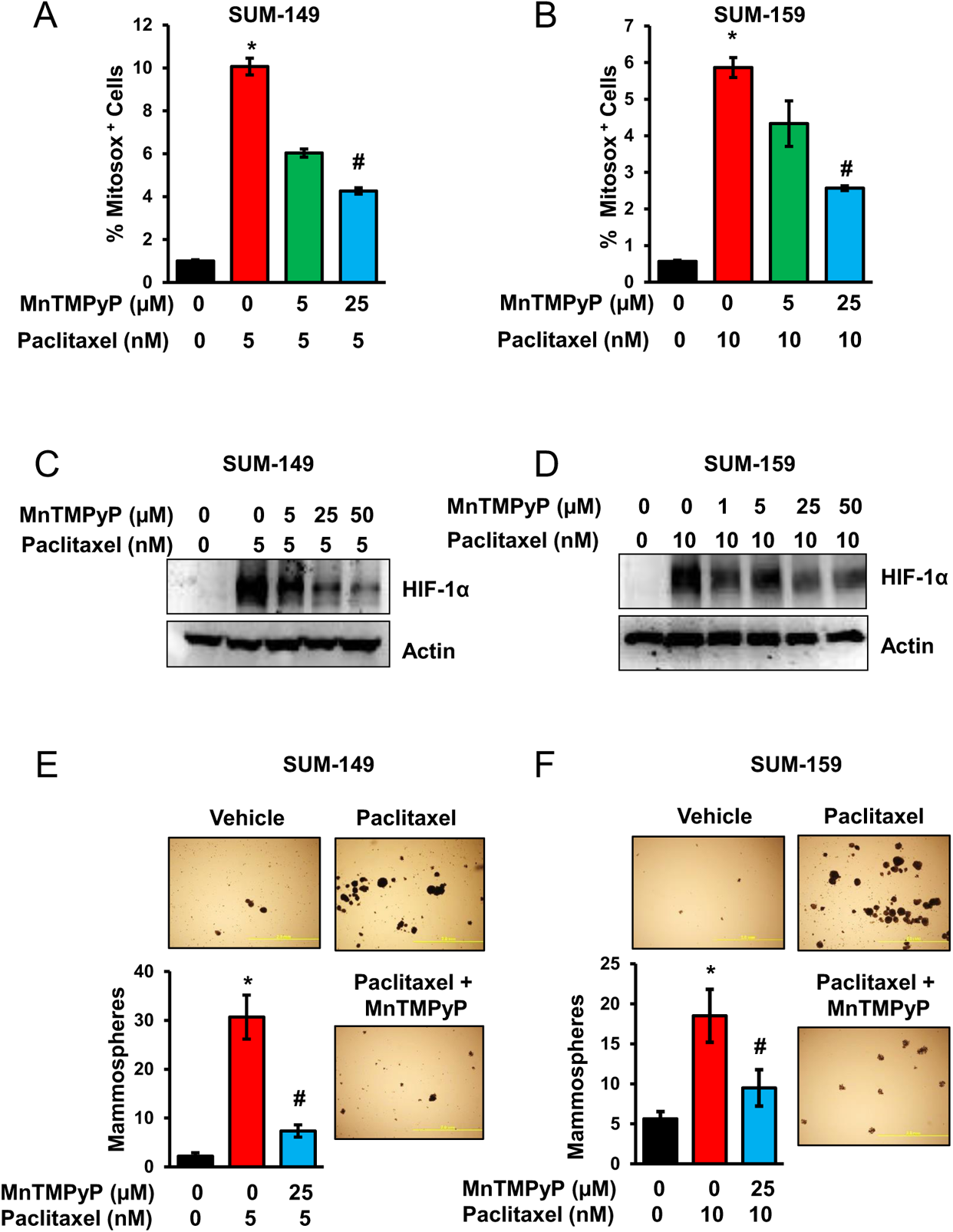
Paclitaxel induces ROS leading to increased levels of HIF-1α and BCSCs. (***A*** and ***B***) Cells were treated as indicated. After four days, Mitosox Red-positive cells were analyzed by flow cytometry (mean ± SEM; *n* = 3). \**p* < 0.001 compared to vehicle-treated cells, and ^#^*p* < 0.001 compared to Pac-treated cells. (***C*** and ***D***) Cells were treated as indicated and IB assays were performed. (***E*** and ***F***) Cells were treated as indicated and mammosphere assays were performed. \**p* < 0.001 compared to vehicle-treated cells, and ^#^*p* < 0.001 compared to Pac-treated cells.

### HIF inhibitors block MDR1 expression in response to paclitaxel

Treatment of TNBC cells with Pac increased MDR1 mRNA and protein expression, which was blocked by digoxin or acriflavine (Figure 6A-C). Flow sorting of TNBC cells revealed that Pac induced a significant increase in MDR1 mRNA levels in ALDH^−^ cells and an even greater induction in ALDH^+^ cells (Figure 6D-E). Knockdown of either or both HIF-α subunits caused a marked decrease in MDR1 mRNA and protein expression, in either vehicle- or Pac-treated cells (Figure 6F-G), indicating that MDR1 expression is highly dependent on HIF activity. When MDA-MB-231 cells were exposed to Pac in the presence of verapamil, which is a competitive inhibitor of MDR1, BCSC enrichment was inhibited (Figure 6H). The data presented in Figure 6 indicate that HIF-dependent MDR1 expression and activity are required for the enrichment BCSCs in response to Pac.

**Figure 6.**
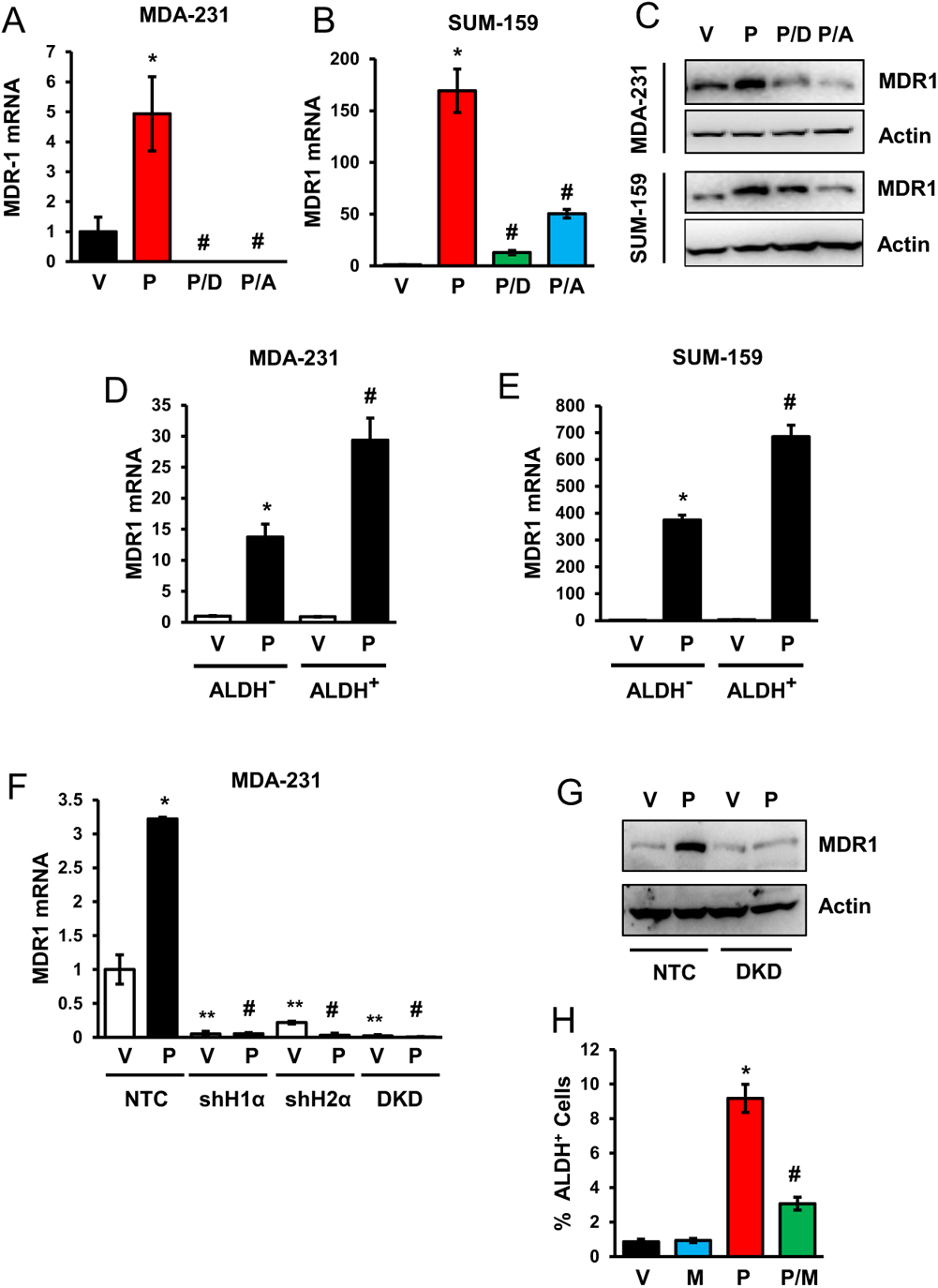
Pac increases expression of MDR1 mRNA and protein. (***A*** and ***B***) Cells were treated with vehicle (V) or Pac, either alone (P) or in combination with digoxin (P/D) or acriflavine (P/A) for four days and RT-qPCR was performed (mean ± SEM; *n* = 3). \**p* < 0.001 compared to V, and ^#^*p* < 0.001 compared to P. (***C***) Cells were treated as described above and IB assays were performed. (***D*** and ***E***) Cells were treated with Pac for four days, sorted into ALDH^−^ and ALDH^+^ populations, and RT-qPCR was performed and normalized to vehicle-treated ALDH^−^ cells. \**p* < 0.001 compared to vehicle-treated ALDH^−^ cells, and ^#^*p* < 0.001 compared to Pac-treated ALDH^−^ cells. (***F***) Subclones were treated with vehicle or Pac and analyzed by RT-qPCR with results normalized to vehicle- treated NTC (mean ± SEM; *n* = 3). \**p* < 0.001 compared to vehicle-treated NTC, and ^#^*p* < 0.001 compared to Pac-treated NTC. (***G***) Subclones were treated with vehicle (V) or Pac (P) for four days and IB assays were performed. (***H***) Cells were treated with vehicle (V), verapamil (M; 50 µM), Pac (P; 10 nM), or both Pac and verapamil (P/M) for four days and ALDH assays were performed (mean ± SEM; *n* = 3). \**p* < 0.001 compared to V, and ^#^*p* < 0.001 compared to P.

### Digoxin blocks the effect of paclitaxel on BCSCs *in vivo*

To determine whether Pac had effects on the BCSC population within tumors, we injected MDA-MB-231 cells into the mammary fat pad (MFP) of immunodeficient *Scid* mice. Once the orthotopic tumors had reached a volume of 200 mm^3^ (designated day 1), the mice received intraperitoneal (IP) injections of saline, digoxin (2 mg/kg every day), Pac (10 mg/kg on days 5 and 10), or both digoxin and Pac. There was no change in body weight of the mice (Figure S4*A*). The combination of Pac and digoxin had a greater inhibitory effect on tumor growth compared to either drug alone (Figure 7*A*). Analysis of tumors harvested on day 12 revealed that Pac increased the abundance of ALDH^+^ cells in the residual tumor (Figure 7*B*). By contrast, digoxin decreased the abundance of ALDH^+^ cells and blocked the increase in ALDH^+^ cells induced by Pac (Figure 7*B*). Pac also increased intratumoral IL-6, IL-8, and MDR1 mRNA levels, which was blocked by co-administration of digoxin (Figure 7*C*).

**Figure 7.**
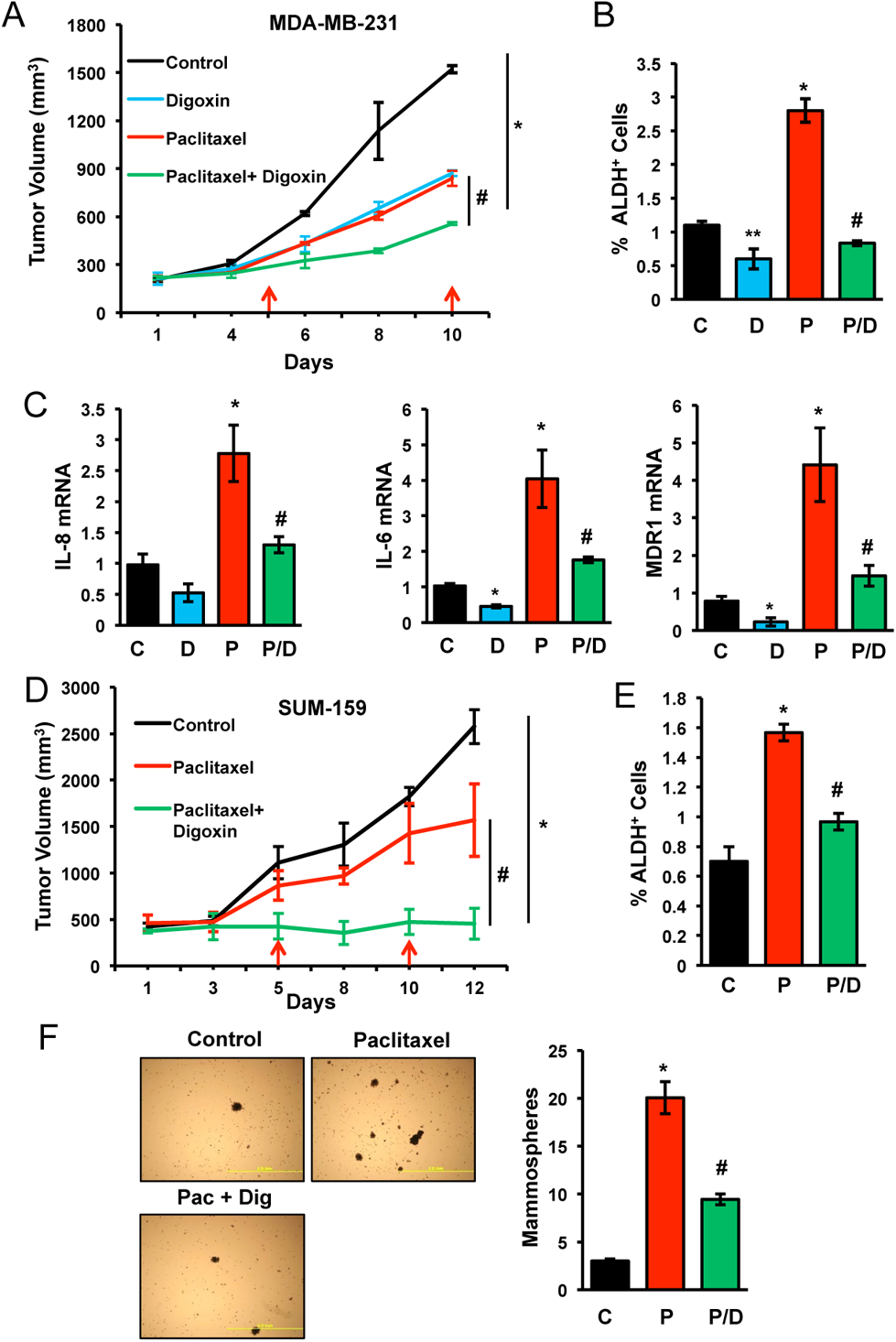
Digoxin blocks BCSC enrichment induced by Pac in vivo. (***A-C***) MDA-MB-231 cells were injected into the mammary fat pad (MFP) of *Scid* mice. When tumors had grown to 200 mm^3^, mice were treated with: saline (Control, C); digoxin (D; 2 mg/kg on days 1-12); Pac (P; 10 mg/kg on days 5 and 10); or Pac and digoxin (P/D). Tumor volumes were measured every 2-3 days (***A***). Tumors were harvested on day 12 for ALDH (***B***) and RT-qPCR (***C***) assays. Data are shown as mean ± SEM (n = 3). \**p* < 0.001 compared to C, and ^#^*p* < 0.001 compared to P. (***D-F***) SUM-159 cells were injected subcutaneously into *Nude* mice. When tumors had grown to 500 mm^3^, mice were treated as described above. Tumor volume was determined every 2-3 days (***D***). Tumors were harvested on day 12 for ALDH (***E***) or mammosphere (***F***) assays. Data are mean ± SEM (n = 3). \**p* < 0.01 compared to C, and ^#^*p* < 0.01 compared to P. Photomicrographs of mammospheres are shown (scale bar: 2 mm).

To validate these findings in a different mouse BC model, we implanted SUM-159 cells subcutaneously into *Nude* mice and started treatment when the tumors grew to 500 mm^3^. Co-administration of Pac and digoxin impaired tumor growth as compared to treatment with saline (control) or Pac alone (Figure 7*D*). Pac increased the percentage of ALDH^+^ cells and digoxin blocked this effect (Figure 7*E*). Pac also increased the abundance of mammosphere-forming cells and digoxin inhibited this effect (Figure 7*F*). None of the treatments caused significant weight loss (Figure S4*B*). The results presented in Figure 7 indicate that combination therapy of TNBC with digoxin blocks Pac-induced expression of IL-6, IL-8 and MDR-1, and inhibits BCSC enrichment *in vivo*.

### Digoxin inhibits BCSC enrichment induced by gemcitabine

To investigate the effect of another cytotoxic chemotherapy drug, we treated MDA-MB-231 and SUM-159 cells with gemcitabine (Gem), which is a nucleoside analog that inhibits DNA synthesis, a mechanism of action distinct from that of Pac, which binds to microtubules and interferes with their disassembly during cytokinesis. HIF-1α expression increased when TNBC cells were exposed to Gem at IC_50_ (10 nM for SUM-159; 20 nM for MDA-MB-231) for four days (Figure 8A-B). Gem also increased the abundance of ALDH^+^ (Figure 8*C*) and mammosphere-forming (Figure 8*D* and Figure S5*A*) cells, and this induction of BCSCs was inhibited by digoxin. Gem increased expression of IL-6 and MDR1, which was blocked by treatment with digoxin or acriflavine (Figure 8E-F and Figure S5B-C). In contrast to Pac, Gem did not increase IL-8 levels in MDA-MB-231 cells (Figure 8*G*). However, treatment with neutralizing antibodies demonstrated that IL-8 signaling was required for BCSC maintenance in the absence of Gem and that signaling mediated by IL-6 and IL-8 was required for the observed enrichment of BCSCs in response to Gem (Figure 8H).

**Figure 8.**
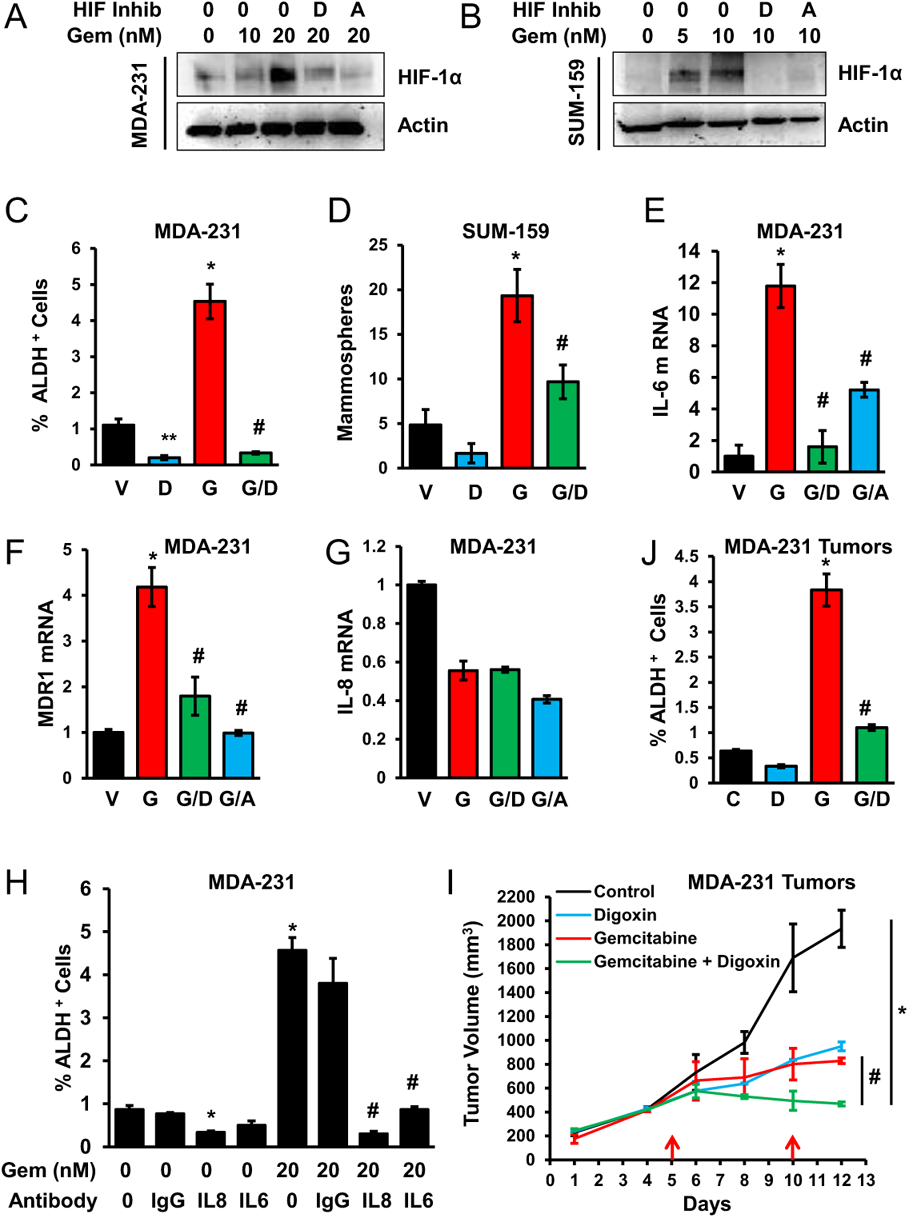
Gem increases BCSC abundance. (***A*** and ***B***) Cells were exposed to gemcitabine (Gem) either alone or in combination with digoxin (D) or acriflavine (A) for four days and IB assays were performed. (***C*** and ***D***) Cells were treated with vehicle (V), digoxin (D), Gem (G; 20 nM for MDA-MB-231 and 10 nM for SUM-159), or Gem and digoxin (G/D) for four days and subjected to ALDH (***C***) and mammosphere (***D***) assays (mean ± SEM; *n* = 3). \**p* < 0.001 compared to V, and ^#^*p* < 0.001 compared to G. (***E-G***) Cells were treated with vehicle (V) or Gem, either alone (*G*) or in combination with digoxin (G/D) or acriflavine (G/A) and RT-qPCR assays were performed (mean ± SEM; n = 3). \**p* < 0.001 compared to V, and ^#^*p* < 0.001 compared to G. (***H***) Cells were treated with vehicle or Gem, either alone or in combination with IgG or neutralizing antibody against IL-6 or IL-8 (500 ng/ml). After four days, ALDH^+^ cells were quantified (mean ± SEM; *n* = 3). \**p* < 0.001 compared to vehicle, and ^#^*p* < 0.001 compared to Gem. (***I*** and ***J***) Cells were injected into the MFP of female *Scid* mice. When tumors had grown to 200 mm^3^, the mice were treated with: saline control; digoxin (2 mg/kg on days 1-12), Gem (20 mg/kg on days 5 and 10); or Gem and digoxin. Tumor volumes were measured every 2 to 3 days (***I***). Tumors were harvested on day 12 and ALDH^+^ cells were quantified (***J***; mean ± SEM; *n* = 3). \**p* < 0.001 compared to control and ^#^*p* < 0.001 compared to Gem.

Next, we injected MDA-MB-231 cells into the MFP of *Scid* mice. When the tumor volume reached 200 mm^3^, mice received IP injections of saline, digoxin (2 mg/kg every day), Gem (20 mg/kg on days 5 and 10), or the combination of digoxin and Gem. Treatment with Gem and digoxin inhibited tumor growth to a greater extent than either drug alone (Figure 8I) without causing weight loss (Figure S4*C*). Gem increased the abundance of ALDH^+^ cells in the residual tumor but digoxin blocked this effect (Figure 8J). The data presented in Figure 8 and Figure S5 indicate that treatment of TNBC cells with digoxin inhibits Gem-induced HIF-1α, IL-6, IL-8 and MDR1 expression, and BCSC enrichment, both in vitro and in vivo.

### Digoxin and gemcitabine combination therapy enables tumor eradication

Compared to patients with BCs that express ER, PR, or HER2, TNBC patients have the greatest risk of relapse within 1–3 years, indicating failure of the initial therapeutic regimen to eradicate the cancer (10, 11). To investigate the conditions required for tumor eradication, MDA-MB-231 cells were injected into *Scid* mice and when the orthotopic tumors became palpable, the mice received IP injections of saline, Gem (20 mg/kg on days 5, 10, 15, 20, and 25), or the combination of Gem and digoxin (2 mg/kg daily). Although treatment with Gem significantly decreased the tumor volume, rapid growth resumed as soon as treatment was discontinued. In contrast, combination therapy eradicated the tumor and prevented any immediate relapse after the discontinuation of Gem treatment (Figure 9).

**Figure 9.**
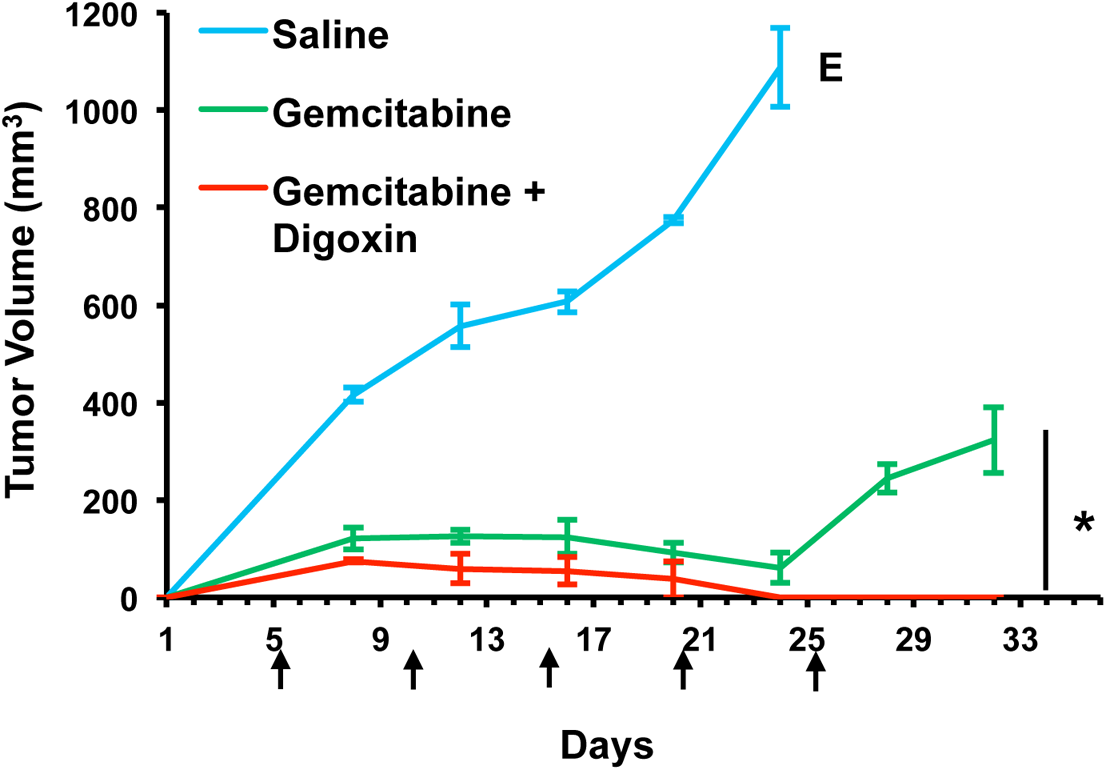
Combined treatment with Gem and digoxin results in tumor eradication. MDA-MB-231 cells were implanted into the MFP of *Scid* mice. When a tumor was palpable (designated day 1), the mice were treated with intraperitoneal injection of: saline; Gem (20 mg/kg on days 5, 10, 15, 20 and 25 [arrows]); or Gem and digoxin (2 mg/kg on days 1-25). Tumor volumes were determined every 2 to 3 days and mean ± SEM (n = 3) values are shown. Saline-treated mice were euthanized on day 24 when tumors exceeded 1000 mm^3^ (E). The experiment was terminated on day 32, seven days after the last dose of Gem. \**p* < 0.001.

### Chemotherapy induces HIF-1α expression and BCSC enrichment in other breast cancer subtypes

To determine whether treatment with Pac induces BCSC enrichment in BC subtypes other than Basal/TNBCs, we exposed MCF-7 (Luminal A subtype/ER^+^PR^+^) and HCC-1954 (HER2^+^) cells to Pac for four days at IC_50_. Pac induced HIF-1α and HIF-2α expression, which was inhibited by treatment with digoxin or acriflavine, in MCF-7 cells (Fig. S6*A*). Pac increased the abundance of ALDH^+^ cells, whereas digoxin decreased the abundance of ALDH^+^ cells, and digoxin inhibited the Pac-induced increase in the abundance of ALDH^+^ cells (Figure S6B). When HCC-1954 cells were exposed to Pac (Figure S6*C*) or Gem (Figure S6*E*) for four days, there was a dose dependent induction of HIF-1α expression, which was inhibited by digoxin or acriflavine. However, there was no enrichment of ALDH^+^ cells after treatment with Pac (Figure S6D) or Gem (Figure S6*F*). The MCF-7 results indicate that chemotherapy-induced BCSC enrichment is not restricted to TNBC cells, whereas the HCC-1954 data suggest that increased HIF-1α expression is not sufficient to increase BCSC specification in this BC cell line.

### HIF target gene expression predicts mortality of chemotherapy-treated BC patients

To explore the relevance of HIF activity with regard to the survival of BC patients, we utilized a HIF-1 signature comprised of 16 genes (*PLOD1, VEGFA, LOX, P4HA2, NDRG1, SLC2A1, ERO1L, ADM, LDHA, PGK1, ANGPTL4, SLC2A3, CA9, HIF1A, IL6,* and *IL8*), which are expressed in human BC cell lines in a HIF-1-dependent manner. This HIF-1 signature was compared to the PAM50 signature, which consists of 50 genes, the expression of which divides human breast cancer specimens into 5 groups (Basal, HER2-enriched, Luminal B, Luminal A, and Normal) (45, 46), with most TNBCs in the Basal subtype (12). Comparison of these two signatures in 1,160 BCs (47) showed enrichment of the HIF-1 signature in the Basal and, to a lesser extent, HER2-enriched subtypes (Figure S7). When HIF-1α mRNA levels were examined in 3,458 BCs (48), levels greater than the median were associated with decreased overall survival (Figure 10*A*), with an even larger survival difference when only Basal BCs were analyzed (Figure 10*B*). Use of the HIF-1 signature resulted in an even more significant difference in patient survival (Figure 10*C*), particularly in Basal subtype BCs (Figure 10*D*). Among BC patients treated only with chemotherapy, the HIF-1 signature were associated with decreased survival (Figure 10*E*), especially among patients with Basal BCs (Figure 10*F*). These clinical correlations are consistent with our *in vitro* and *in vivo* mechanistic studies demonstrating HIF-dependent enrichment of BCSCs in TNBCs after chemotherapy.

**Figure 10.**
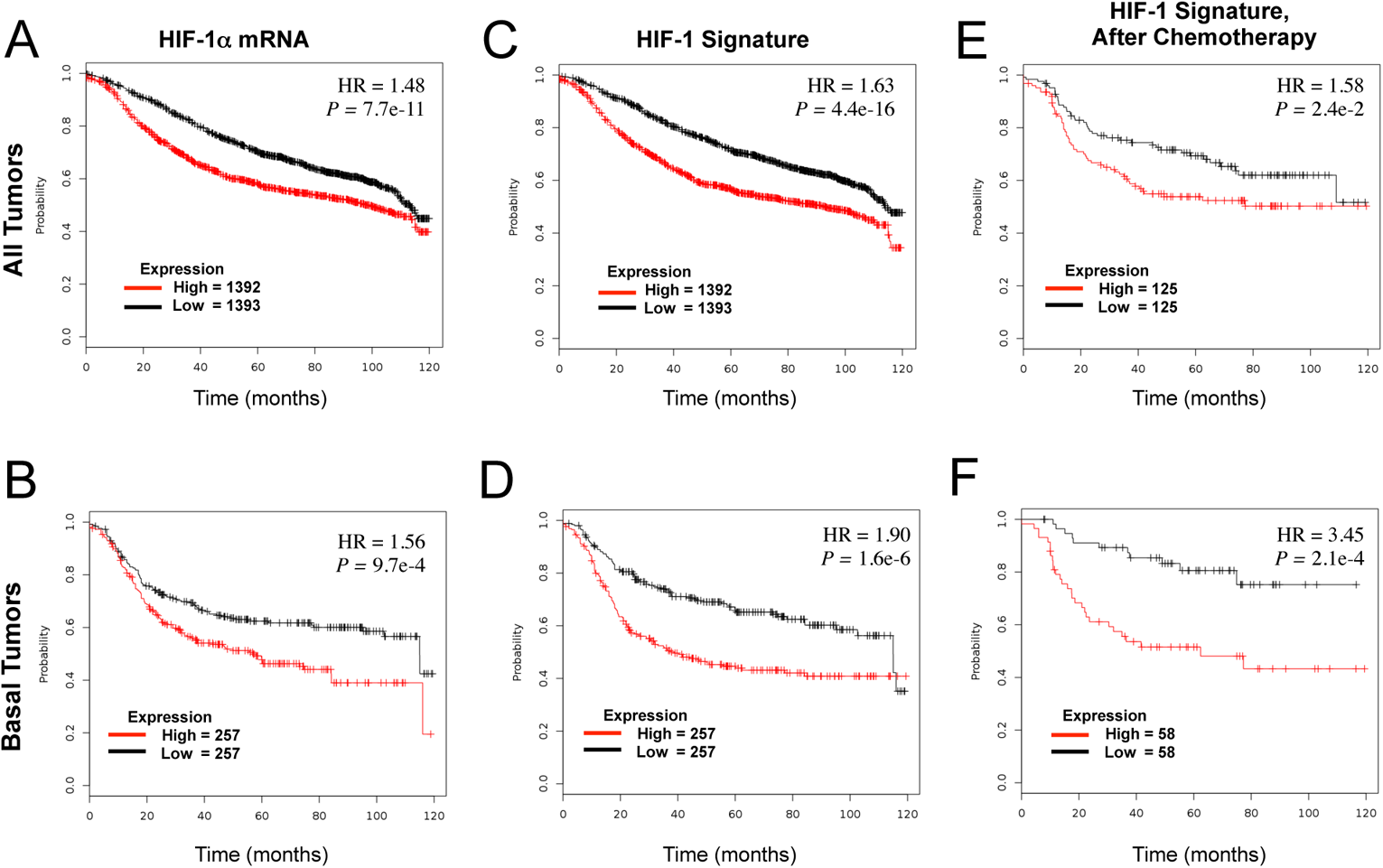
HIF-1α and HIF-1 target gene mRNA expression in primary BCs predict clinical outcome. Kaplan–Meier plots of overall survival (120 months) are shown for 2,785 BC patients, who were stratified according to expression in the primary tumor of HIF-1α mRNA (***A*** and ***B***) or a 16-gene HIF-1 signature (***C-F***), which was greater (red) or less (black) than the median. The *p* value (log-rank test) and hazard ratio (HR) for each comparison are shown. Analyses were performed that included all BC types (***A*** and ***C***) or only Basal-type BC (***B*** and ***D***). Within these groups, the subgroup who received only chemotherapy (**E** and **F**) was analyzed.

## Discussion

By contrast to advances in the treatment of BCs expressing hormone receptors or HER2, effective therapy for women with TNBC remains an unmet clinical need. A large and growing body of experimental data indicates that BCSC eradication is required to achieve a durable remission, but BCSCs are more resistant to chemotherapy that bulk cancer cells (4–6). Previous work has revealed that HIF-1α expression is induced by doxorubicin (24) and that HIF-1α-deficient cells have increased sensitivity to the cytotoxic effects of carboplatin and etoposide (15). In this paper, we have reported that exposure of TNBC cells to clinically relevant concentrations of Pac or Gem increases ROS levels, which induce HIF-1α and HIF-2α mRNA and protein expression, leading to expression of MDR-1, IL-8 and/or IL-6, and to increased abundance of BCSCs, both *in vitro* and *in vivo* (Figure 11). IL-6 and IL-8 have been shown to play essential roles in TNBC tumorigenesis and are associated with increased BC patient mortality (37). IL-6 (33, 35) and IL-8 (6, 34) are required for the maintenance of BCSCs and their resistance to cytotoxic chemotherapy, particularly in TNBCs.

**Figure 11.**
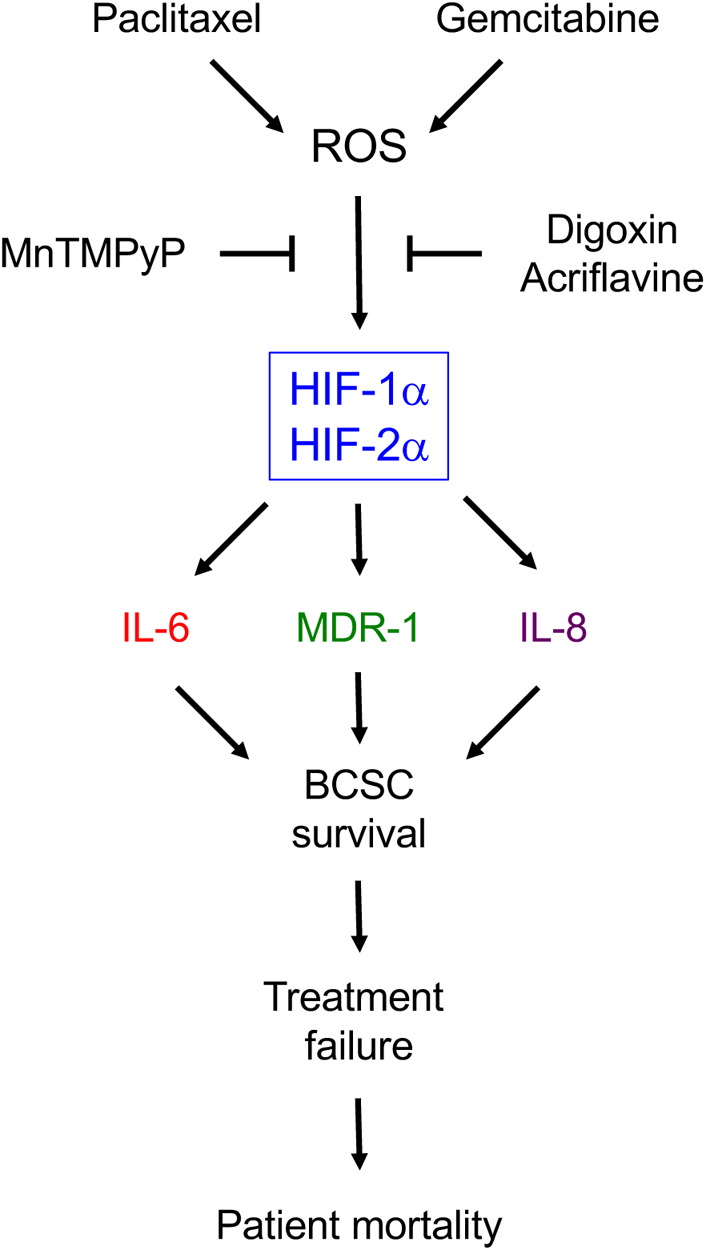
Treatment of TNBC with Pac or Gem induces ROS, leading to HIF-1α and HIF-2α expression, and subsequent expression of IL-6, IL-8, and MDR1, which promote BCSC survival. This counter-therapeutic response is a critical determinant of treatment failure and patient mortality.

It should be noted that treatment with Pac or Gem induced HIF-1α and HIF-2α expression under non-hypoxic conditions and that both HIF-1α and HIF-2α were required for maximal BCSC expansion in response to Pac, whereas only HIF-1α was required to increase the percentage of BCSCs under hypoxic conditions (22), which implies distinct mechanisms leading to BCSC augmentation in response to chemotherapy vs hypoxia that require further study. Pac and Gem induced IL-6 expression and BCSC augmentation, but only Pac increased IL-8 levels, although basal IL-8 expression was required for the increase in BCSC abundance that was induced by Pac. In agreement with these findings, inhibitors of CXCR1, which is the cognate receptor for IL-8, have been shown to decrease BCSC abundance (34).

The increase in IL-8 expression and BCSC abundance induced by Pac was reported to require TGF-β signaling to SMAD2/4 (6), whereas IL-6 expression was reported to require STAT3 activation in order to induce BCSC enrichment (36). However, we found that HIF deficiency in TNBCs abrogated the increased levels of IL-6, IL-8, and BCSCs that were induced by Pac, yet phosphorylation of SMAD2 and STAT3 was still induced. These results indicated that Pac increased SMAD2 and STAT3 activity in a HIF-independent manner but activation of these transcription factors did not lead to BCSC enrichment in the absence of HIF activity. We also found that expression of HIF-1α but not IL-8, was induced in Gem-treated MDA-MB-231 cells, and that Pac or Gem treatment of HCC-1954 cells induced HIF-1α expression but not BCSC enrichment, indicating that HIF-1α expression is not sufficient for IL-8 expression or BCSC enrichment. Taken together, our data and prior studies suggest that HIFs, SMADs, and STAT3 all contribute to the BCSC augmentation that is induced by Pac or Gem. From a translational point of view, it should be noted that genetic or pharmacologic inhibition of HIF activity is sufficient to block BCSC augmentation in response to cytotoxic chemotherapy.

IL-6 and IL-8 signaling augment BCSCs by blocking apoptosis and by stimulating BCSC specification in response to cytotoxic chemotherapy. MDR1 expression acts to maintain viability by pumping Pac or Gem out of cancer cells. Treatment of TNBC cells with either Pac or Gem increased MDR1 expression in a HIF-dependent manner in ALDH^−^ and ALDH^+^ TNBC cells, providing a mechanism for increased resistance of BCSCs to chemotherapy relative to bulk cancer cells. IL-8 signaling also contributes to this differential resistance because CXCR1 expression is restricted to BCSCs (34).

Combination therapy with digoxin blocked the stimulatory effects of Pac or Gem on IL-6, IL-8, and MDR1 expression, leading to decreased tumor growth and blocking BCSC enrichment in two different mouse models of TNBC. Treatment of TNBC-tumor-bearing mice with the HSP90 inhibitor ganetespib induced HIF-1α (but not HIF-2α) degradation, blocked primary tumor growth, and decreased the percentage of ALDH^+^ cells in the residual tumor (49), providing independent evidence that HIF inhibitors prevent BCSC enrichment. HIF inhibition by methylselenocysteine sensitized head and neck squamous cell carcinoma xenografts to irinotecan (50), suggesting that HIF activity promotes cancer stem cell enrichment in response to cytotoxic chemotherapy in other cancer types and may be blocked by other HIF inhibitors.

The clinical relevance of these results is underscored by reports of an association between: (a) ALDH expression and ER^−^/Basal subtype BC (51); (b) HIF-1α and ALDH^+^ cells in BCs, both before and after chemotherapy (51); (c) ALDH^+^ BC cells and resistance to treatment with Pac and epirubicin (52); and (d) ALDH^+^ cells after chemotherapy and decreased disease-free survival (51, 53). Our mechanistic studies complement these clinical correlations and provide support for clinical trials combining HIF inhibitors and chemotherapy in women with TNBC. Several drugs that inhibit HIF activity (54, 55) could be considered for such trials.

## Methods

### Cell Culture

Cells were cultured in high-glucose DMEM (MDA-MB-231 and MCF-7), DMEM/F12 with hydrocortisone and insulin (SUM-159 and SUM-149), or RPMI-1640 (HCC-1954), which was supplemented with 10% fetal bovine serum (FBS) and 1% penicillin/streptomycin. MDA-MB-231 knockdown subclones were cultured in the presence of 0.5 μg/ml of puromycin. Cells were maintained at 37°C in a 5% CO_2_ and 95% air incubator (20% O_2_). Pac, digoxin, acriflavine and Gem were purchased from Sigma-Aldrich and dissolved in DMSO. MnTMPyP was purchased from Calbiochem and dissolved in deionized water.

### RT-qPCR

RNA was extracted using TRIzol (Invitrogen) and treated with DNase I (Ambion). cDNA was synthesized using a 1-μg aliquot of RNA and the iScript cDNA Synthesis system (Bio-Rad). qPCR was performed using human-specific primers (Table S1) and iQ SYBR Green Supermix (Bio-Rad), with annealing temperature optimized by gradient PCR. The expression (E) of each mRNA relative to 18*S* rRNA was calculated based on the cycle threshold (Ct): E = 2^−Δ(ΔCt)^, in which ΔCt = Ct_target_ – Ct_18S_ and Δ(ΔCt) = ΔCt_treatment_ – ΔCt_control_.

### IB Assays

Cells were lysed in RIPA lysis buffer (ThermoFisher). Blots were probed with antibodies against HIF-1α, HIF-2α, IL-6, IL-8, MDR1, phospho-SMAD2, phospho-STAT3, SMAD2, and STAT3 (Novus Biologicals), followed by HRP-conjugated anti-rabbit (Roche) or anti-mouse (Santa Cruz) secondary antibodies and signal was detected using ECL Plus (GE Healthcare). Blots were stripped and re-probed with anti-actin antibody (Santa Cruz).

### Luciferase Assay

MDA-MB-231 cells were seeded overnight and transfected with reporter plasmids pSV-RL and p2.1 (56) using PolyJet (SignaGen). The media was changed after 6 hours and allowed to recover overnight. Cells were treated with vehicle or 10 nM Pac for 4 days, lysed, and dual luciferase activities (Promega) were measured in a multiwell luminescence reader (Perkin–Elmer Life Science).

### ALDH Assay

The Aldefluor assay (StemCell Technologies) was utilized to identify ALDH^+^ cells. Trypsinized cells or minced tumor tissue was digested with type 1 collagenase (1 mg/ml; Sigma) at 37 °C for 30 minutes and passed through a 70-μm strainer. One million viable cells were suspended in assay buffer containing BODIPY aminoacetaldehyde (1 μM) and incubated for 45 minutes at 37 °C. As a negative control, an aliquot of cells was treated with the ALDH inhibitor diethylaminobenzaldehyde (50 mM). Samples were passed through a 35-µm strainer and analyzed by flow cytometry (FACScalibur, BD Biosciences).

### Mammosphere Assay

Cells were seeded on ultra-low attachment plates (Corning) at a density of 5×10^3^ cells/ml in Complete MammoCult Medium (StemCell Technologies). After seven days, cells were photographed under an Olympus TH4-100 microscope with ×4 apochromat objective lens. Mammospheres with area > 500 pixels were counted in images of 3 fields per well in triplicate wells and the mean number of mammospheres per field was calculated. For secondary mammosphere formation, primary mammospheres were trypsinized, plated at 5×10^3^ cells/ml, incubated for seven days and analyzed as described above.

### MitoSOX Staining

Cells were incubated in 5 μM MitoSOX Red (Molecular Probes) at 37 °C for 45 minutes in phosphate-buffered saline (PBS) with 5% FBS, followed by rinsing with PBS. Stained cells were filtered and subjected to flow cytometry (FACScan, BD Bioscience). The gain and amplifier settings were held constant for all samples.

### Mouse Studies

Protocols were in accordance with the NIH *Guide for the Care and Use of Laboratory Animals* and were approved by the Johns Hopkins University Animal Care and Use Committee. Female five- to seven-week-old *Scid* and *Nude* mice (NCI) were implanted with MDA-MB-231 or SUM-159 cells by MFP or subcutaneous injection, respectively. Pac, Gem, digoxin, and saline for injection were obtained from the research pharmacy of the Johns Hopkins Hospital. Cells were suspended at 2×10^7^ cells/ml in a 1:1 solution of PBS and Matrigel (Corning). Primary tumors were measured in 3 dimensions (*a*, *b*, *c*), and volume (V) was determined based on the formula: V = *abc* × 0.52.

### Statistical Analysis

Data are expressed as mean ± SEM. Differences between 2 groups and multiple groups were analyzed by Student’s *t* test and ANOVA, respectively. *P* values < 0.05 were considered significant. For the HIF-1 signature, the Breast Invasive Carcinoma gene expression dataset of 1,162 patients was analyzed (47). Kaplan-Meier curves were generated from a BC dataset containing mRNA expression and overall survival data on 2,785 patients (48). The log rank test was performed to determine whether observed differences between groups were significant. HIF-1 signature expression was analyzed according to BC subtype using the GOBO database (57).

## ACKNOWLEDGMENTS

We thank Vered Stearns of the Kimmel Cancer Center at Johns Hopkins for helpful discussions. We are grateful to Karen Padgett of Novus Biologicals for providing IgG and antibodies against HIF-2α, IL-6, IL-8, MDR1, p-SMAD2, SMAD2, p-STAT3, STAT3, and p-IκBα. This work was supported by: Breast Cancer Research Program Impact Award W81XWH-12-1-0464 from the Department of Defense; Cigarette Restitution Fund Research Grant FH-B33-CRF from the State of Maryland Department of Health and Mental Hygiene; the Cindy Rosencrans Fund for Triple Negative Breast Cancer; and the WTFC. D.M.G. was supported by National Cancer Institute grant K99-CA181352. G.L.S. is an American Cancer Society Research Professor and the C. Michael Armstrong Professor at Johns Hopkins University School of Medicine.

## Supplemental Figures

**Figure S1.**
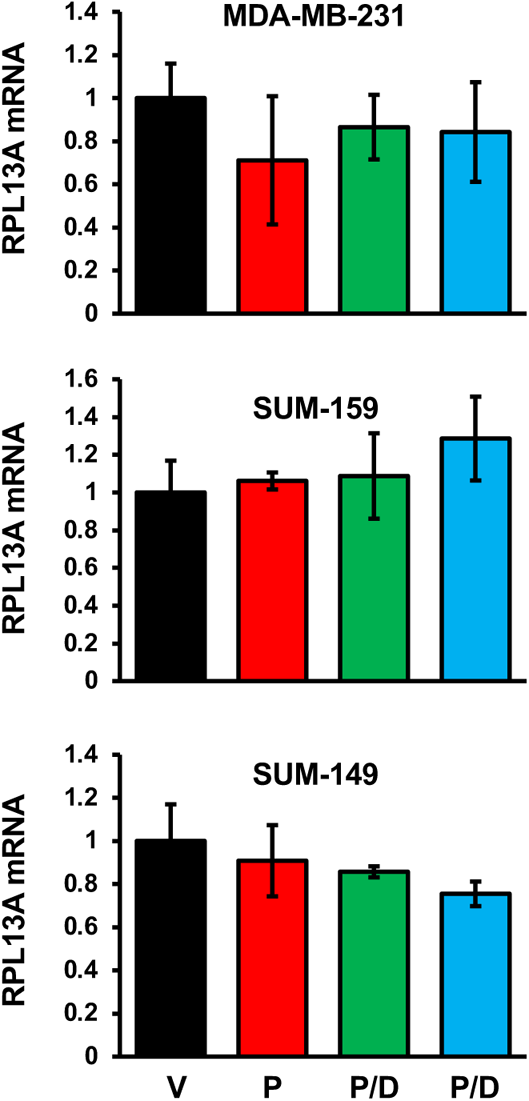
Analysis of RPL13A mRNA expression in TNBC cells. Cells were treated with vehicle or Pac, either alone (P) or in combination with digoxin (P/D) or acriflavine (P/A) for four days. RT-qPCR analyses were performed to quantify RPL13A mRNA levels relative to 18S rRNA in the same sample and normalized to vehicle treated cells (mean ± SEM; *n* = 3).

**Figure S2.**
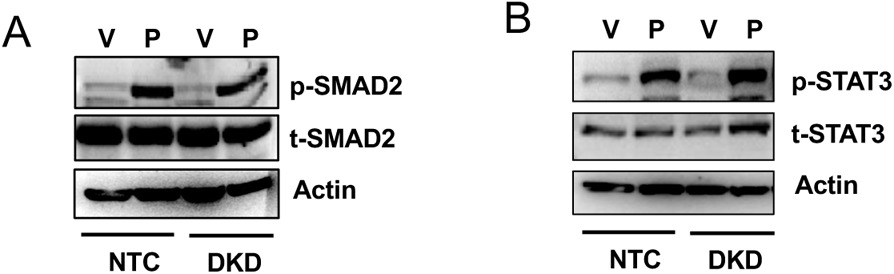
Phosphorylation of SMAD2 and STAT3 in Pac-treated MDA-MB-231 subclones. NTC and DKD subclones were treated with either vehicle or Pac for four days and IB assays were performed using antibodies specific for phosphorylated (p-) and total (t-) SMAD2 (***A***) or STAT3 (***B***).

**Figure S3.**
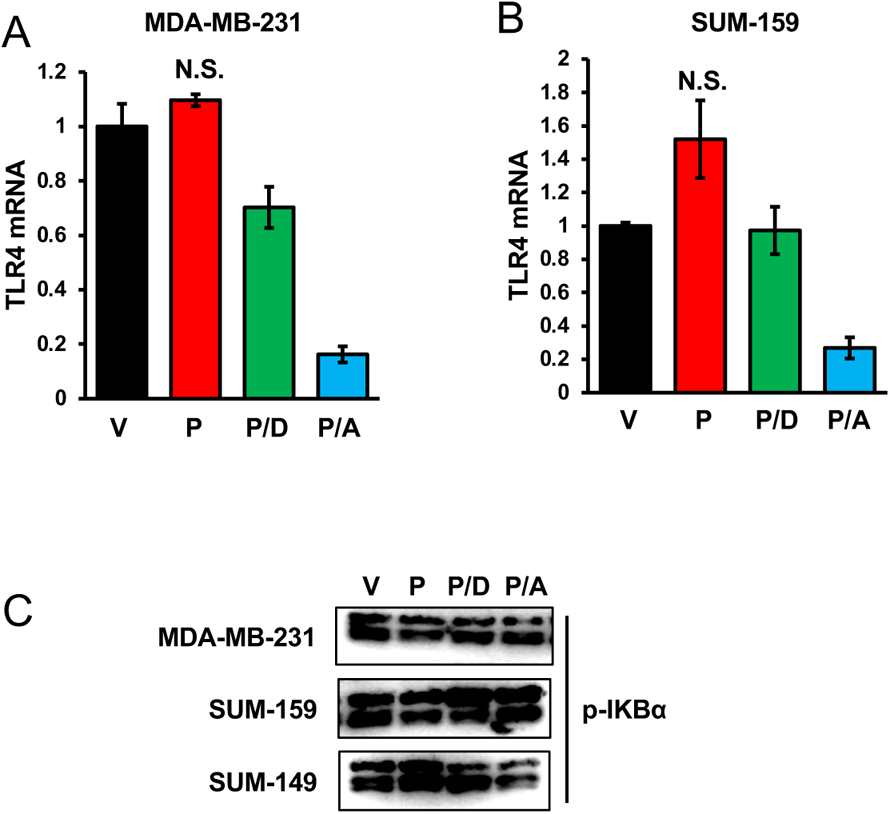
Analysis of TLR-4 expression and IκB phosphorylation in Pac-treated cells. (***A*** and ***B***) TNBC cells were treated with vehicle (V) or Pac, either alone (P) or in combination with digoxin (P/D) or acriflavine (P/A) for four days. TLR-4 mRNA levels were quantified relative to 18S rRNA in the same sample and normalized to V (mean ± SEM; *n* = 3). N.S., not significant. (***C***) TNBC cells were treated as described above and IB assays were performed using antibody against phospho-IκBα.

**Figure S4.**
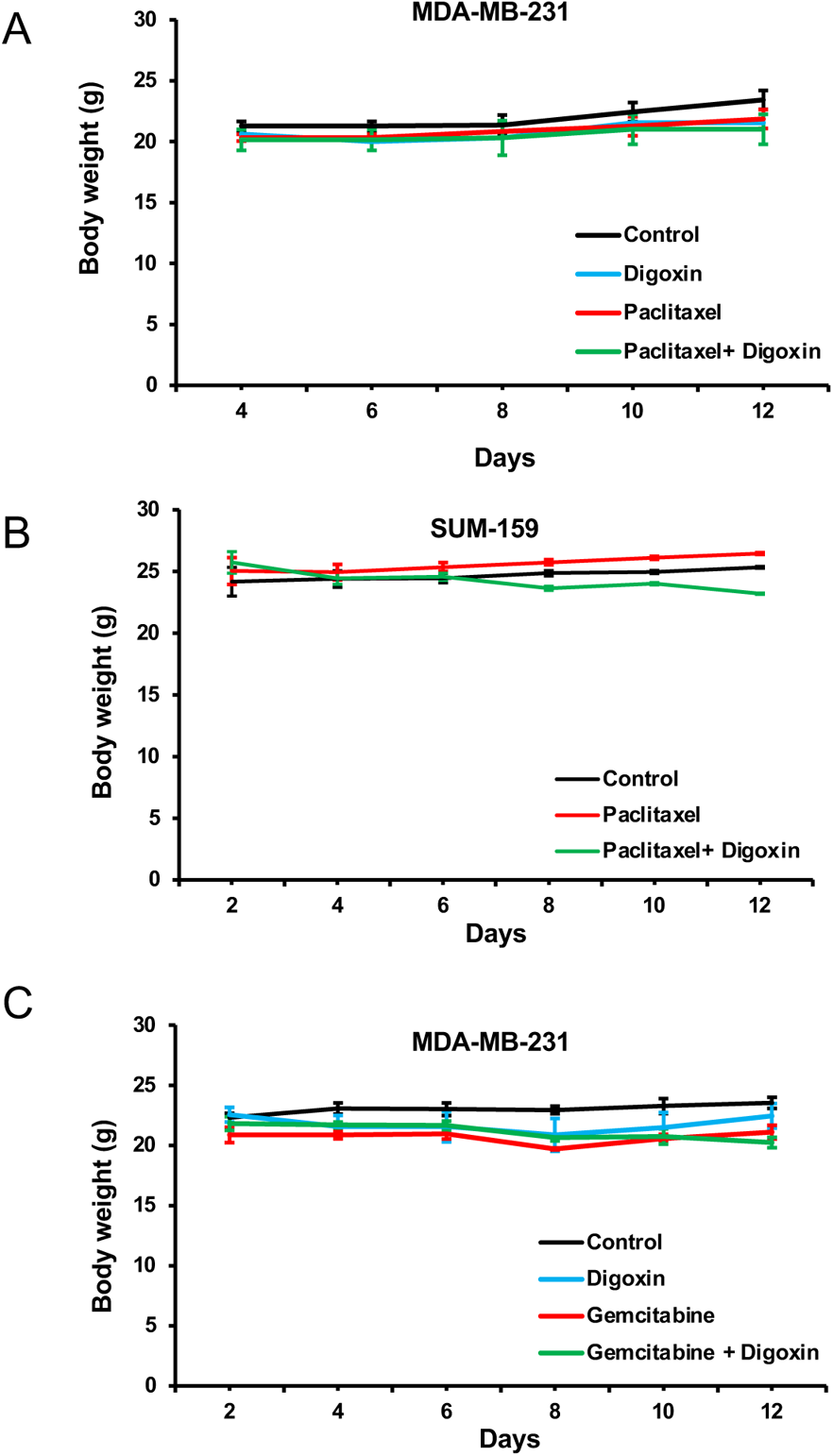
Body weights of tumor-bearing mice treated with chemotherapy or digoxin. *Scid* mice received MFP implantation of MDA-MB-231 cells (***A*** and ***C***) and *Nude* mice received subcutaneous implantation of SUM-159 cells (***B***). Body weights were measured every 2 to 3 days during the treatment with saline (control), digoxin (days 1-12), chemotherapy (Pac or Gem, days 5 and 10) or digoxin plus Pac or Gem.

**Figure S5.**
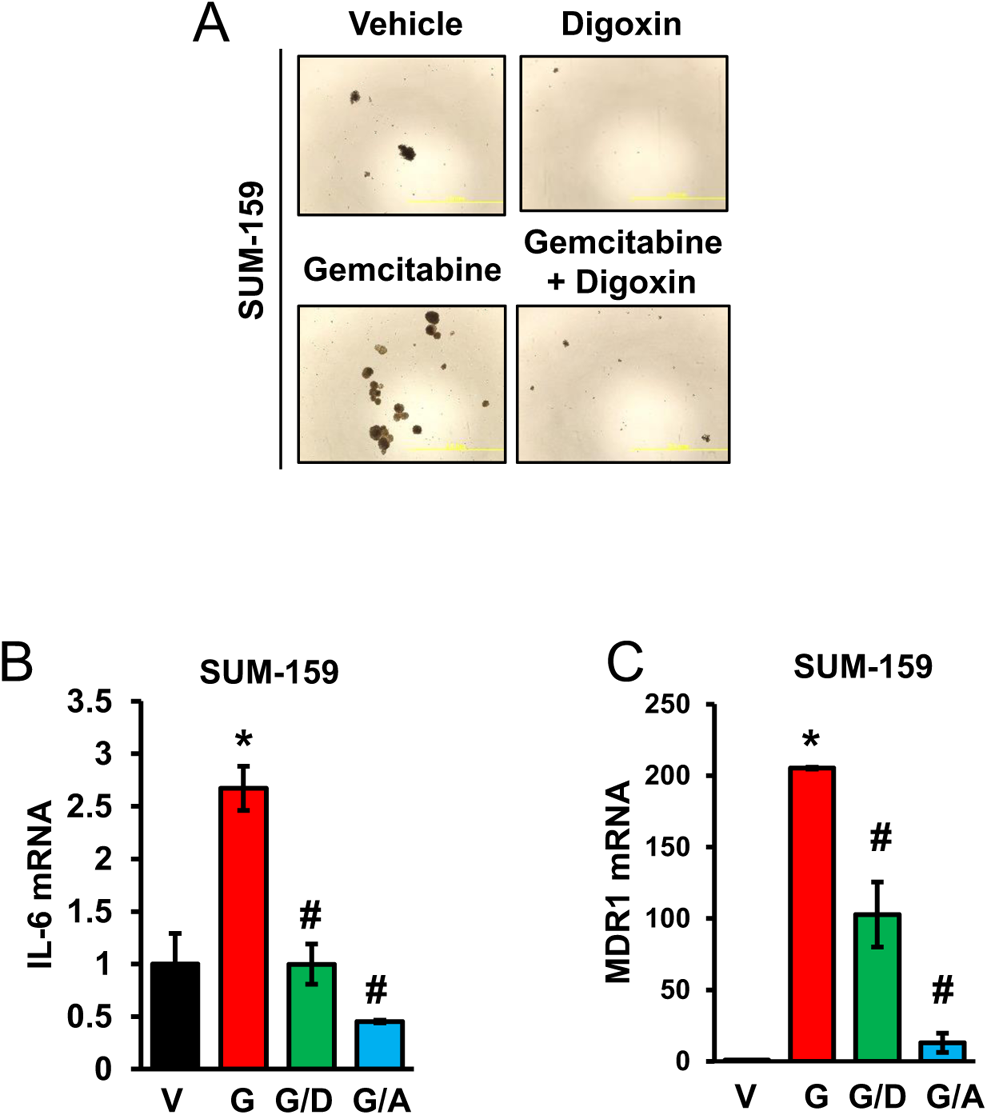
Gem increased BCSCs and IL-6 and MDR-1 mRNA levels in a HIF-dependent manner. (***A***) Cells were treated with vehicle, 100 nM digoxin, 20 nM Gem, or Gem plus digoxin for four days and mammosphere assays were performed. Photomicrographs of mammospheres are shown (scale bar: 2 mm). (***B*** and ***C***) Cells were treated with vehicle (V) or Gem, either alone (G) or in combination with digoxin (G/D) or acriflavine (G/A). IL-6 (***B***) and MDR1 (***C***) mRNA levels were determined relative to 18S rRNA, with expression normalized to vehicle-treated cells (mean ± SEM; *n* = 3). \**p* < 0.001 compared to V, and ^#^*p* < 0.001 compared to G.

**Figure S6.**
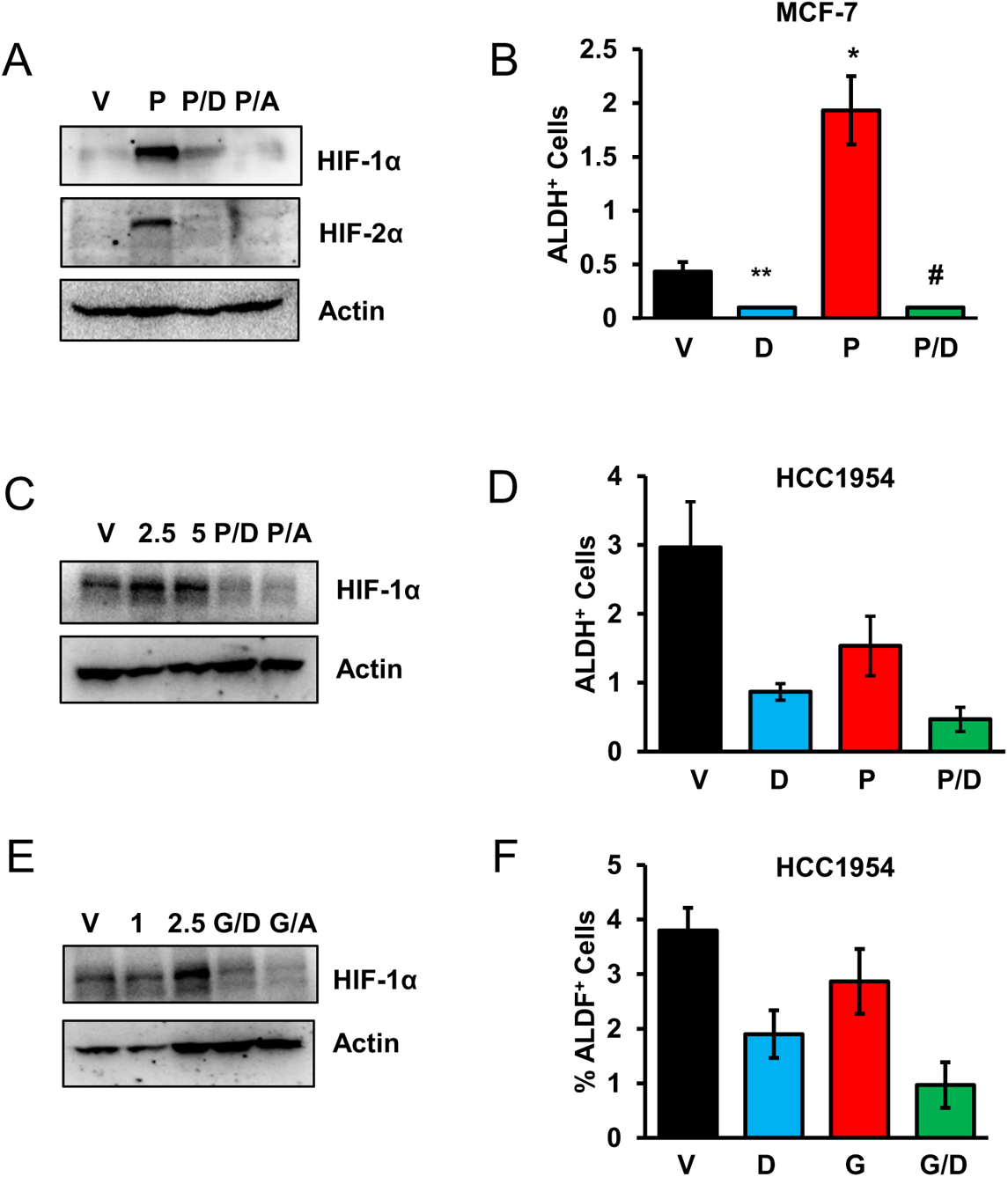
Effect of Pac or Gem treatment on non-basal-type BC cells. (***A*** and ***B***) ER^+^ MCF-7 cells were treated with vehicle (V) or 10 nM paclitaxel, either alone (P) or in combination with digoxin (P/D) or acriflavine (P/A) for four days and IB (***A***) or ALDH (***B***) assays were performed (mean ± SEM; *n* = 3). \**p* < 0.001 compared to V, and ^#^*p* < 0.001 compared to P. (***C***) HER2^+^ HCC-1954 cells were treated with vehicle (V), 2.5 or 5 nM Pac alone, or 2.5 nM Pac in combination with digoxin (P/D) or acriflavine (P/A) and IB assays were performed. (***D***) Cells were treated with vehicle (V), digoxin (D), Pac (P; 2.5 nM), or both Pac and digoxin (P/D) and ALDH assays were performed (mean ± SEM; *n* = 3). (***E***) Cells were treated with vehicle (V), 1 or 2.5 nM Gem alone, or 2.5 nM Gem plus digoxin (G/D) or acriflavine (G/A) and IB assays were performed. (***F***) Cells were treated with vehicle (V), digoxin (D), Gem (G; 2.5 nM), or Gem plus digoxin (G/D) and ALDH assays were performed (mean ± SEM; *n* = 3).

**Figure S7.**
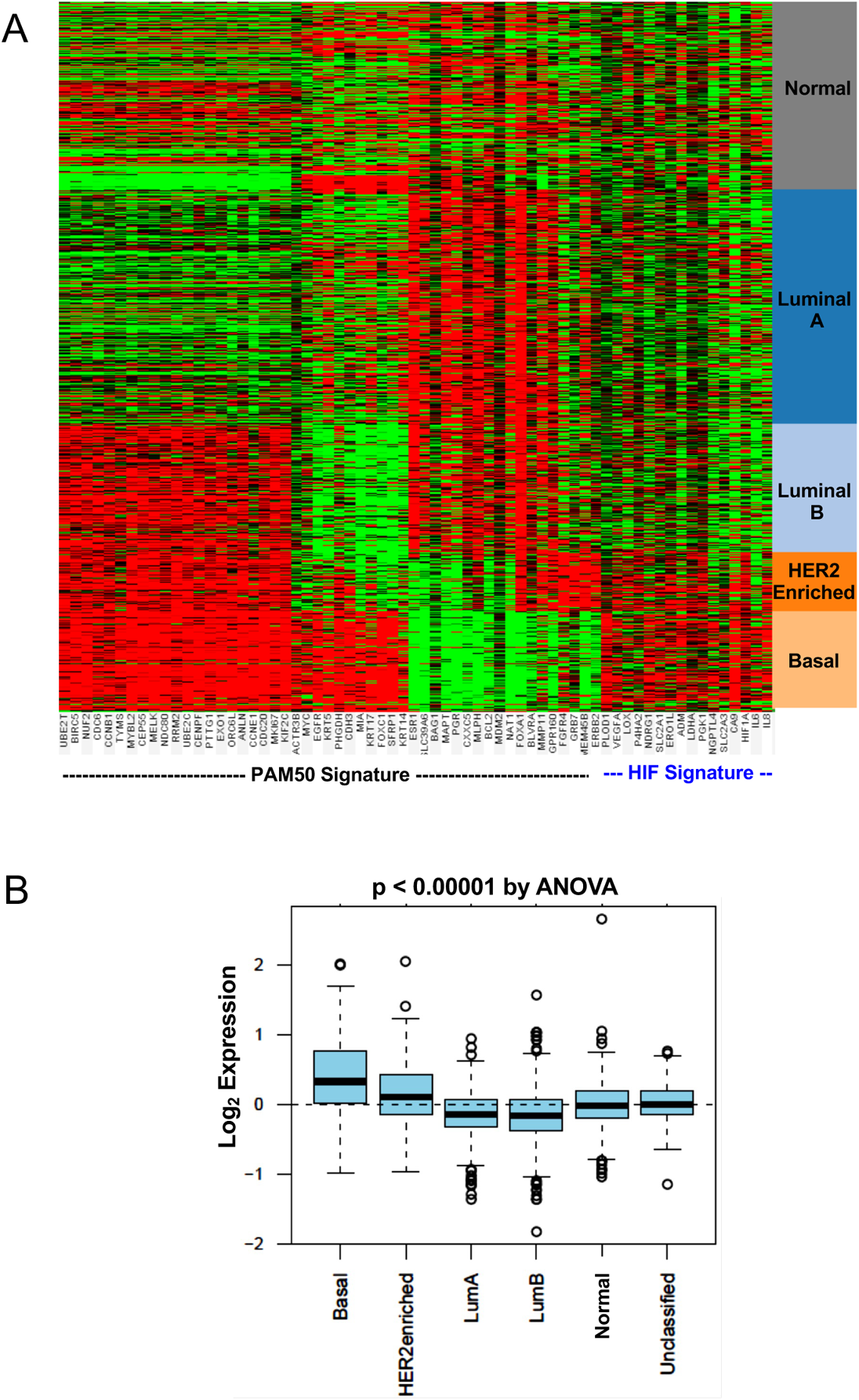
Expression of HIF-1 target genes in primary BCs. (***A***) Expression of PAM50 (46) and HIF signatures in 1,160 primary BCs was used to classify tumors into those with gene expression that was greater (red) or less (green) than the median. (***B***) HIF signature was significantly associated with Basal subtype in 1,160 BCs using the GOBO database (57). Lum, Luminal.

**Table S1.**
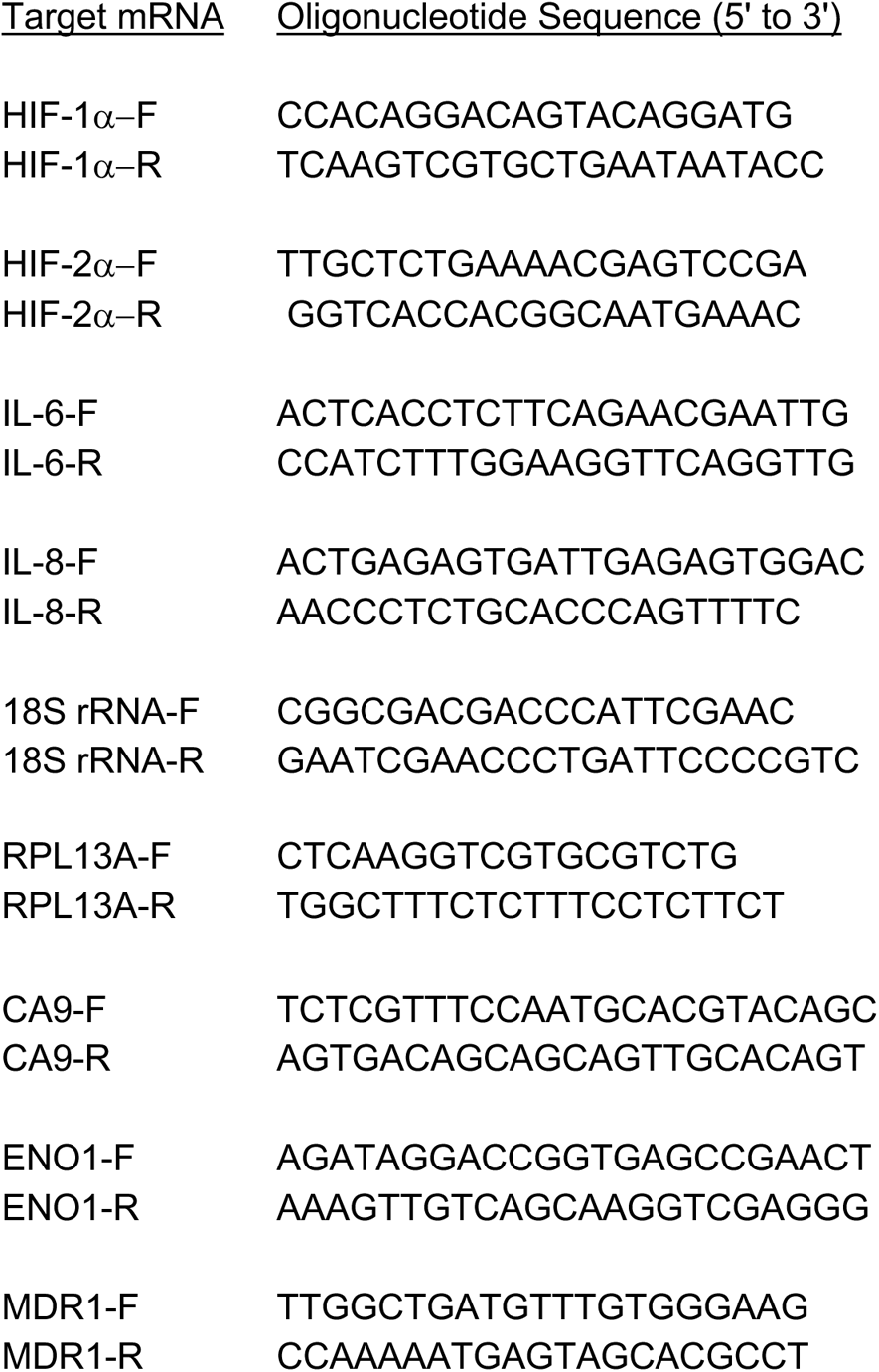
Sequence of forward (F) and reverse (R) primers used for RT-qPCR.

